# Large-scale exome sequencing study implicates both developmental and functional changes in the neurobiology of autism

**DOI:** 10.1101/484113

**Authors:** F. Kyle Satterstrom, Jack A. Kosmicki, Jiebiao Wang, Michael S. Breen, Silvia De Rubeis, Joon-Yong An, Minshi Peng, Ryan Collins, Jakob Grove, Lambertus Klei, Christine Stevens, Jennifer Reichert, Maureen S. Mulhern, Mykyta Artomov, Sherif Gerges, Brooke Sheppard, Xinyi Xu, Aparna Bhaduri, Utku Norman, Harrison Brand, Grace Schwartz, Rachel Nguyen, Elizabeth E. Guerrero, Caroline Dias, Branko Aleksic, Richard Anney, Mafalda Barbosa, Somer Bishop, Alfredo Brusco, Jonas Bybjerg-Grauholm, Angel Carracedo, Marcus C.Y. Chan, Andreas G. Chiocchetti, Brian H. Y. Chung, Hilary Coon, Michael L. Cuccaro, Aurora Currò, Bernardo Dalla Bernardina, Ryan Doan, Enrico Domenici, Shan Dong, Chiara Fallerini, Montserrat Fernández-Prieto, Giovanni Battista Ferrero, Christine M. Freitag, Menachem Fromer, J. Jay Gargus, Daniel Geschwind, Elisa Giorgio, Javier González-Peñas, Stephen Guter, Danielle Halpern, Emily Hansen-Kiss, Xin He, Gail E. Herman, Irva Hertz-Picciotto, David M. Hougaard, Christina M. Hultman, Iuliana Ionita-Laza, Suma Jacob, Jesslyn Jamison, Astanand Jugessur, Miia Kaartinen, Gun Peggy Knudsen, Alexander Kolevzon, Itaru Kushima, So Lun Lee, Terho Lehtimäki, Elaine T. Lim, Carla Lintas, W. Ian Lipkin, Diego Lopergolo, Fátima Lopes, Yunin Ludena, Patricia Maciel, Per Magnus, Behrang Mahjani, Nell Maltman, Dara S. Manoach, Gal Meiri, Idan Menashe, Judith Miller, Nancy Minshew, Eduarda Montenegro M. de Souza, Danielle Moreira, Eric M. Morrow, Ole Mors, Preben Bo Mortensen, Matthew Mosconi, Pierandrea Muglia, Benjamin Neale, Merete Nordentoft, Norio Ozaki, Aarno Palotie, Mara Parellada, Maria Rita Passos-Bueno, Margaret Pericak-Vance, Antonio Persico, Isaac Pessah, Kaija Puura, Abraham Reichenberg, Alessandra Renieri, Evelise Riberi, Elise B. Robinson, Kaitlin E. Samocha, Sven Sandin, Susan L. Santangelo, Gerry Schellenberg, Stephen W. Scherer, Sabine Schlitt, Rebecca Schmidt, Lauren Schmitt, Isabela Maya W. Silva, Tarjinder Singh, Paige M. Siper, Moyra Smith, Gabriela Soares, Camilla Stoltenberg, Pål Suren, Ezra Susser, John Sweeney, Peter Szatmari, Lara Tang, Flora Tassone, Karoline Teufel, Elisabetta Trabetti, Maria del Pilar Trelles, Christopher Walsh, Lauren A. Weiss, Thomas Werge, Donna Werling, Emilie M. Wigdor, Emma Wilkinson, Jeremy A. Willsey, Tim Yu, Mullin H.C. Yu, Ryan Yuen, Elaine Zachi, iPSYCH consortium, Catalina Betancur, Edwin H. Cook, Louise Gallagher, Michael Gill, James S. Sutcliffe, Audrey Thurm, Michael E. Zwick, Anders D. Børglum, Matthew W. State, A. Ercument Cicek, Michael E. Talkowski, David J. Cutler, Bernie Devlin, Stephan J. Sanders, Kathryn Roeder, Mark J. Daly, Joseph D. Buxbaum

**Affiliations:** Stanley Center for Psychiatric Research, Broad Institute, Cambridge, Massachusetts, USA; Analytic and Translational Genetics Unit, Department of Medicine, Massachusetts General Hospital, Boston, Massachusetts, USA; Program in Medical and Population Genetics, Broad Institute of MIT and Harvard, Cambridge, Massachusetts, USA; Harvard Medical School, Boston, Massachusetts, USA; Center for Genomic Medicine, Department of Medicine, Massachusetts General Hospital, Boston, Massachusetts, USA; Department of Statistics, Carnegie Mellon University, Pittsburgh, Pennsylvania, USA; Seaver Autism Center for Research and Treatment, Icahn School of Medicine at Mount Sinai, New York, New York, USA; Department of Psychiatry, Icahn School of Medicine at Mount Sinai, New York, New York, USA; The Mindich Child Health and Development Institute, Icahn School of Medicine at Mount Sinai, New York, New York, USA; Department of Psychiatry, UCSF Weill Institute for Neurosciences, University of California San Francisco, San Francisco, California, USA; Program in Bioinformatics and Integrative Genomics, Harvard Medical School, Boston, Massachusetts, USA; The Lundbeck Foundation Initiative for Integrative Psychiatric Research, iPSYCH, Denmark; Center for Genomics and Personalized Medicine, Aarhus, Denmark; Department of Biomedicine - Human Genetics, Aarhus University, Aarhus, Denmark; Department of Psychiatry, University of Pittsburgh School of Medicine, Pittsburgh, Pennsylvania, USA; Department of Neurology, University of California San Francisco, San Francisco, California, USA; The Eli and Edythe Broad Center of Regeneration Medicine and Stem Cell Research, University of California San Francisco, San Francisco, California, USA; Computer Engineering Department, Bilkent University, Ankara, Turkey; Center for Autism Research and Translation, University of California Irvine, Irvine, California, USA; MIND (Medical Investigation of Neurodevelopmental Disorders) Institute, University of California Davis, Davis, California, USA; Division of Genetics, Boston Children’s Hospital, Boston, Massachusetts, USA; Division of Developmental Medicine, Boston Children’s Hospital, Boston, Massachusetts, USA; Department of Psychiatry, Graduate School of Medicine, Nagoya University, Nagoya, Japan; Division of Psychological Medicine & Clinical Neurosciences, MRC Centre for Neuropsychiatric Genetics & Genomics, Cardiff University, Cardiff, United Kingdom; Department of Medical Sciences, University of Torino, Turin, Italy; Medical Genetics Unit, Città della Salute e della Scienza Hospital, Turin, Italy; Center for Neonatal Screening, Department for Congenital Disorders, Statens Serum Institut, Copenhagen, Denmark; Grupo de Medicina Xenómica, Centro de Investigaciónen en Red de Enfermedades Raras (CIBERER), Universidade de Santiago de Compostela, Santiago de Compostela, Spain; Fundación Pública Galega de Medicina Xenómica, Servicio Galego de Saúde (SERGAS), Santiago de Compostela, Spain; Department of Pediatrics & Adolescent Medicine, Duchess of Kent Children’s Hospital, The University of Hong Kong, Hong Kong Special Administrative Region, China; Department of Child and Adolescent Psychiatry, Psychosomatics and Psychotherapy, Goethe University Frankfurt, Frankfurt, Germany; Department of Internal Medicine, University of Utah, Salt Lake City, Utah, USA; Department of Psychiatry, University of Utah, Salt Lake City, Utah, USA; John P Hussman Institute for Human Genomics, University of Miami, Miami, USA; Medical Genetics, University of Siena, Siena, Italy; Child Neuropsychiatry Department, Department of Surgical Sciences, Dentistry, Gynecology and Pediatrics. University of Verona, Verona, Italy; Centre for Integrative Biology, University of Trento, Trento, Italy; Neurogenetics Group, Instituto de Investigación Sanitaria de Santiago (IDIS-SERGAS), Santiago de Compostela, Spain; Department of Public Health and Pediatrics, University of Torino, Turin, Italy; Department of Human Genetics, David Geffen School of Medicine, University of California Los Angeles, Los Angeles, California, USA; Center for Autism Research and Treatment and Program in Neurobehavioral Genetics, Semel Institute, David Geffen School of Medicine, University of California Los Angeles, Los Angeles, California, USA; Department of Child and Adolescent Psychiatry, Hospital General Universitario Gregorio Marañón, IiSGM, CIBERSAM, School of Medicine Complutense University, Madrid, Spain; Institute for Juvenile Research, Department of Psychiatry, University of Illinois at Chicago, Chicago, Illinois, USA; The Research Institute at Nationwide Children’s Hospital, Columbus, Ohio, USA; Department of Human Genetics, University of Chicago, Chicago, Illinois, USA; Department of Medical Epidemiology and Biostatistics, Karolinska Institutet, Stockholm, Sweden; Department of Biostatistics, Columbia University, New York, New York, USA; Norwegian Institute of Public Health, Oslo, Norway; Department of Child Psychiatry, University of Tampere and Tampere University Hospital, Tampere, Finland; Department of Pediatrics, Icahn School of Medicine at Mount Sinai, New York, New York, USA; Department of Clinical Chemistry, Fimlab Laboratories and Finnish Cardiovascular Research Center-Tampere, Faculty of Medicine and Life Sciences, University of Tampere, Tampere, Finland; Campus Bio-medico, Rome, Italy; Life and Health Sciences Research Institute, School of Medicine, University of Minho, Campus de Gualtar, Braga, Portugal; Department of Psychiatry, Massachusetts General Hospital and Harvard Medical School, Boston, Massachusetts, USA; Pre-School Psychiatry Unit, Soroka University Medical Center, Beer Sheva, Israel; Department of Public Health, Ben-Gurion University of the Negev, Beer-Sheva, Israel; Centro de Pesquisas sobre o Genoma Humano e Células tronco, Departamento de Genética e Biologia Evolutiva, Instituto de Biociências, Universidade de São Paulo, São Paulo, Brazil; Department of Molecular Biology, Cell Biology and Biochemistry and Department of Psychiatry and Human Behavior, Brown University, Providence, Rhode Island, USA; Psychosis Research Unit, Aarhus University Hospital, Risskov, Denmark; National Centre for Register-Based Research, Aarhus University, Aarhus, Denmark; Centre for Integrated Register-based Research, Aarhus University, Aarhus, Denmark; UCB Pharma, Belgium; Mental Health Services in the Capital Region of Denmark, Mental Health Center Copenhagen, University of Copenhagen, Copenhagen, Denmark; Institute for Molecular Medicine Finland (FIMM), University of Helsinki, Helsinki, Finland; Psychiatric & Neurodevelopmental Genetics Unit, Department of Psychiatry, Massachusetts General Hospital, Boston, Massachusetts, USA; Interdepartmental Program “Autism 0-90”, Gaetano Martino University Hospital, University of Messina, Messina, Italy; Mafalda Luce Center for Pervasive Developmental Disorders, Milan, Italy; Department of Environmental Medicine and Public Health, Icahn School of Medicine at Mount Sinai, New York, New York, USA; Genetica Medica, Azienda Ospedaliera Universitaria Senese, Siena, Italy; The Wellcome Trust Sanger Institute, Cambridge, United Kingdom; Maine Medical Center and Maine Medical Center Research Institute, Portland, Maine, USA; Department of Psychiatry, Tufts University School of Medicine, Boston, Massachusetts, USA; Department of Pathology and Laboratory Medicine, University of Pennsylvania School of Medicine, Philadelphia, Pennsylvania, USA; Program in Genetics and Genome Biology, The Centre for Applied Genomics, The Hospital for Sick Children, Toronto, Ontario, Canada; McLaughlin Centre, University of Toronto, Toronto, Ontario, Canada; Center for Medical Genetics Dr. Jacinto Magalhães, National Health Institute Dr. Ricardo Jorge, Porto, Portugal; New York State Psychiatric Institute, New York, New York, USA; Department of Epidemiology, Mailman School of Public Health, Columbia University, Epidemiology, Mailman School of Public Health, Columbia University, New York, New York, USA; Department of Psychiatry and Behavioural Neurosciences, Offord Centre for Child Studies, McMaster University, Hamilton, Ontario, Canada; Department of Biochemistry and Molecular Medicine, University of California Davis, School of Medicine, Davis, CA, USA; Department of Neurosciences, Biomedicine and Movement Sciences, Section of Biology and Genetics, University of Verona, Verona, Italy; Institute of Biological Psychiatry, MHC Sct. Hans, Mental Health Services Copenhagen, Roskilde, Denmark; Department of Clinical Medicine, University of Copenhagen, Copenhagen, Denmark; Sorbonne Université, INSERM, CNRS, Neuroscience Paris Seine, Institut de Biologie Paris Seine, Paris, France; Department of Psychiatry, School of Medicine, Trinity College Dublin, Dublin, Ireland; Vanderbilt Genetics Institute, Vanderbilt University School of Medicine, Nashville, Tennessee, USA; Department of Molecular Physiology and Biophsysics and Psychiatry, Vanderbilt University School of Medicine, Nashville, Tennessee, USA; National Institute of Mental Health, National Institutes of Health, Bethesda, Maryland, USA; Department of Human Genetics, Emory University School of Medicine, Atlanta, Georgia, USA; Bioinformatics Research Centre, Aarhus University, Aarhus, Denmark; Computational Biology Department, Carnegie Mellon University, Pittsburgh, Pennsylvania, USA; Department of Genetics and Genomic Sciences, Icahn School of Medicine at Mount Sinai, New York, New York, USA; Friedman Brain Institute, Icahn School of Medicine at Mount Sinai, New York, New York, USA; Department of Neuroscience, Icahn School of Medicine at Mount Sinai, New York, New York, USA

## Abstract

We present the largest exome sequencing study of autism spectrum disorder (ASD) to date (n=35,584 total samples, 11,986 with ASD). Using an enhanced Bayesian framework to integrate *de novo* and case-control rare variation, we identify 102 risk genes at a false discovery rate ≤ 0.1. Of these genes, 49 show higher frequencies of disruptive *de novo* variants in individuals ascertained for severe neurodevelopmental delay, while 53 show higher frequencies in individuals ascertained for ASD; comparing ASD cases with mutations in these groups reveals phenotypic differences. Expressed early in brain development, most of the risk genes have roles in regulation of gene expression or neuronal communication (i.e., mutations effect neurodevelopmental and neurophysiological changes), and 13 fall within loci recurrently hit by copy number variants. In human cortex single-cell gene expression data, expression of risk genes is enriched in both excitatory and inhibitory neuronal lineages, consistent with multiple paths to an excitatory/inhibitory imbalance underlying ASD.

## Introduction

Autism spectrum disorder (ASD), a childhood-onset neurodevelopmental condition characterized by deficits in social communication and restricted, repetitive patterns of behavior or interests, affects more than 1% of individuals (Baio et al., 2018). Multiple studies have demonstrated high heritability, much of it due to common variation (Gaugler et al., 2014), although rare inherited and *de novo* variants are major contributors to individual risk (De Rubeis et al., 2014; Iossifov et al., 2014; Sanders et al., 2015). When this rare variation disrupts a gene in individuals with ASD more often than expected by chance, it implicates that gene in risk (He et al., 2013). ASD risk genes, in turn, provide insight into the underpinnings of ASD, both individually (Ben-Shalom et al., 2017; Bernier et al., 2014) and *en masse* (De Rubeis et al., 2014; Ruzzo et al., 2018; Sanders et al., 2015; Willsey et al., 2013). However, fundamental questions about the altered neurodevelopment and altered neurophysiology in ASD—including when it occurs, where, and in what cell types—remain poorly resolved.

Here we present the largest exome sequencing study in ASD to date. Through an international collaborative effort and the willingness of thousands of participating families, we assembled a cohort of 35,584 samples, including 11,986 with ASD. We introduce an enhanced Bayesian analytic framework, which incorporates recently developed gene- and variant-level scores of evolutionary constraint of genetic variation, and we use it to identify 102 ASD-associated genes (FDR ≤ 0.1). Because ASD is often one of a constellation of symptoms of neurodevelopmental delay (NDD), we identify subsets of the 102 ASD-associated genes that have disruptive *de novo* variants more often in NDD-ascertained or ASD-ascertained cohorts. We also consider the cellular function of ASD-associated genes and, by examining extant data from single cells in the developing human cortex, show that their expression is enriched in maturing and mature excitatory and inhibitory neurons from midfetal development onwards, confirm their role in neuronal communication or regulation of gene expression, and show that these functions are separable. Together, these insights form an important step forward in elucidating the neurobiology of ASD.

## Results

### Dataset

We analyzed whole-exome sequence data from 35,584 samples that passed our quality control procedures (STAR Methods). This included 21,219 family-based samples (6,430 ASD cases, 2,179 unaffected controls, and both of their parents) and 14,365 case-control samples (5,556 ASD cases, 8,809 controls) (Fig. S1; Table S1). Of these, 17,462 samples were either newly sequenced by our consortium (6,197 samples: 1,908 probands with parents; 274 ASD cases; 25 controls) or newly incorporated (11,265 samples: 416 probands with parents; 4,811 ASD cases and 5,214 controls from the Danish iPSYCH study (Satterstrom et al., 2018)).

From this cohort, we identified a set of 9,345 rare *de novo* variants in protein-coding exons (allele frequency ≤ 0.1% in our dataset as well as in the non-psychiatric subsets of the reference databases ExAC and gnomAD, with 63% of probands and 59% of unaffected offspring carrying at least one such rare coding *de novo* variant—4,073 out of 6,430 and 1,294 out of 2,179, respectively; Table S2; Fig. S1). For rare inherited and case-control analyses, we included variants with an allele count no greater than five in our dataset and in the non-psychiatric subset of ExAC (Kosmicki et al., 2017; Lek et al., 2016).

### Impact of genetic variants on ASD risk

The differential burden of genetic variants carried by cases versus controls reflects the average liability they impart for ASD. For example, because protein-truncating variants (PTVs, consisting of nonsense, frameshift, and essential splice site variants) show a greater difference in burden between ASD cases and controls than missense variants, their average impact on liability must be larger (He et al., 2013). Recent analyses have shown that measures of functional severity, such as the “probability of loss-of-function intolerance” (pLI) score (Kosmicki et al., 2017; Lek et al., 2016) and the integrated “missense badness, PolyPhen-2, constraint” (MPC) score (Samocha et al., 2017), can further delineate variant classes with higher burden. Therefore, we divided the list of rare autosomal genetic variants into seven tiers of predicted functional severity—three tiers for PTVs by pLI score (≥0.995, 0.5-0.995, 0-0.5), in order of decreasing expected impact; three tiers for missense variants by MPC score (≥2, 1-2, 0-1), also in order of decreasing impact; and a single tier for synonymous variants, expected to have minimal impact. We further divided the variants by their inheritance pattern: *de novo*, inherited, and case-control. Unlike inherited variants, *de novo* mutations are exposed to minimal selective pressure and have the potential to mediate substantial risk to disorders that limit fecundity, including ASD (Power et al., 2013). This expectation is borne out by the substantially higher proportions of all three PTV tiers and the two most severe missense variant tiers in *de novo* variants compared to inherited variants (Fig. 1A).

**Figure 1.**
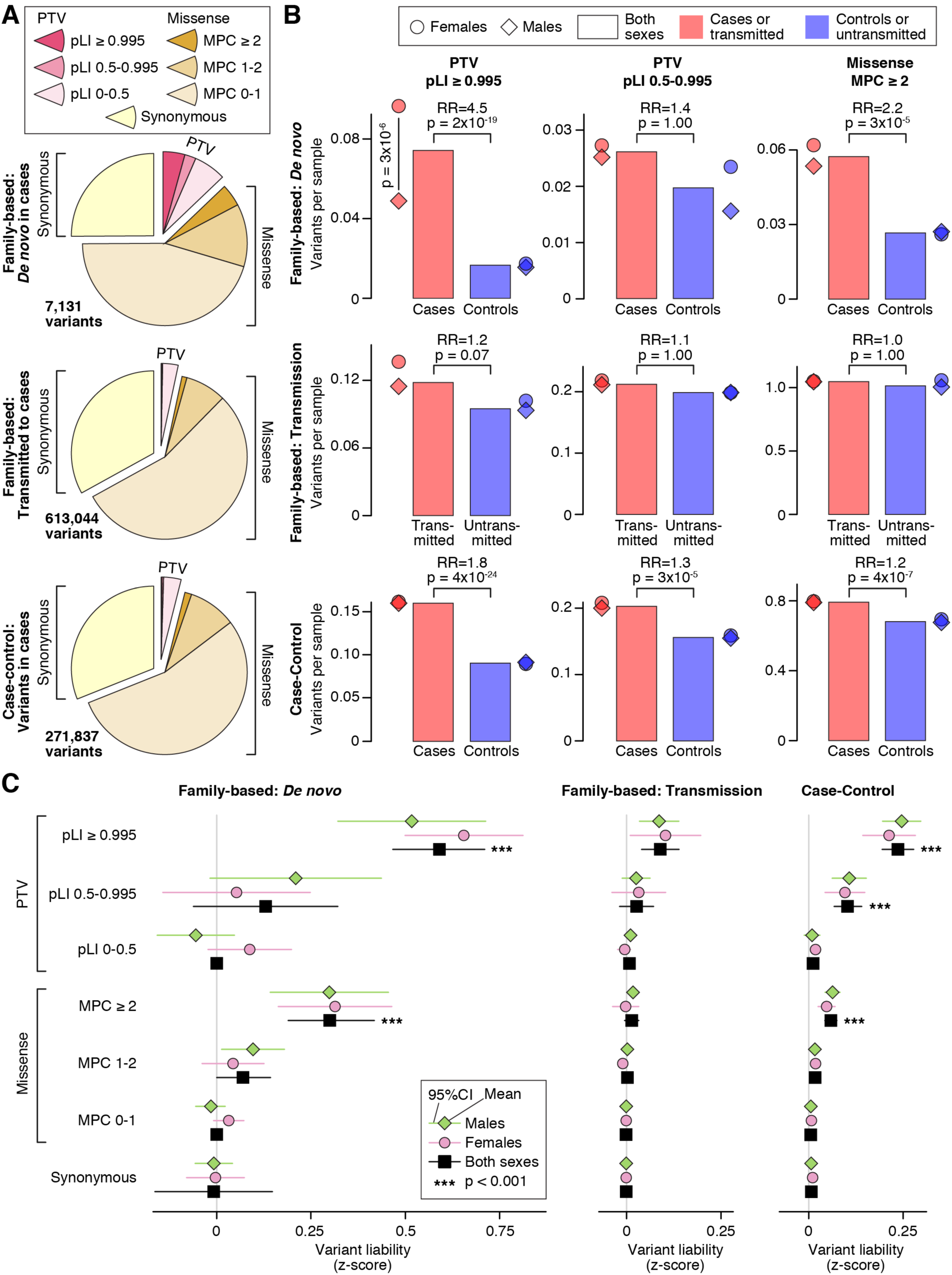
Distribution of rare autosomal protein-coding variants in ASD cases and controls. **A**, The proportion of rare autosomal genetic variants split by predicted functional consequences, represented by color, is displayed for family-based data (split into de novo and inherited variants) and case-control data. PTVs and missense variants are split into three tiers of predicted functional severity, represented by shade, based on the pLI and MPC metrics, respectively. **B**, The relative difference in variant frequency (i.e. burden) between ASD cases and controls (top and bottom) or transmitted and untransmitted parental variants (middle) is shown for the top two tiers of functional severity for PTVs (left and center) and the top tier of functional severity for missense variants (right). Next to the bar plot, the same data are shown divided by sex. **C**, The relative difference in variant frequency shown in ‘B’ is converted to a trait liability z-score, split by the same subsets used in ‘A’. For context, a z-score of 2.18 would shift an individual from the population mean to the top 1.69% of the population (equivalent to an ASD threshold based on 1 in 68 children (Christensen et al., 2016)). No significant difference in liability was observed between males and females for any analysis. Statistical tests: B, C: Binomial Exact Test (BET) for most contrasts; exceptions were “both” and “case-control”, for which Fisher’s method for combining BET p-values for each sex and, for case-control, each population, was used; p-values corrected for 168 tests are shown.

Comparing probands to unaffected siblings, we observe a 3.5-fold enrichment of *de novo* PTVs in the 1,447 autosomal genes with a pLI ≥ 0.995 (366 in 6,430 cases versus 35 in 2,179 controls; 0.057 vs. 0.016 variants per sample (vps); p=4×10^−17^, two-sided Poisson exact test; Fig. 1B). A less pronounced difference is observed for rare inherited PTVs in these genes, with a 1.2-fold enrichment of transmitted versus untransmitted alleles (695 transmitted versus 557 untransmitted in 5,869 parents; 0.12 vs. 0.10 vps; p=0.07, binomial exact test; Fig. 1B). The relative burden in the case-control data falls between the estimates for *de novo* and inherited data in these most severe PTVs, with a 1.8-fold enrichment in cases versus controls (874 in 5,556 cases versus 759 in 8,809 controls; 0.16 vs. 0.09 vps; p=4×10^−24^, binomial exact test; Fig. 1B). Analysis of the middle tier of PTVs (0.5 ≤ pLI < 0.995) shows a similar, but muted, pattern (Fig. 1B), while the lowest tier of PTVs (pLI < 0.5) shows no case enrichment (Table S3).

*De novo* missense variants are observed more frequently than *de novo* PTVs and, *en masse*, they show only marginal enrichment over the rate expected by chance (De Rubeis et al., 2014) (Fig. 1). However, the most severe *de novo* missense variants (MPC ≥ 2) occur at a frequency similar to *de novo* PTVs, and we observe a 2.1-fold case enrichment (354 in 6,430 cases versus 58 in 2,179 controls; 0.055 vs. 0.027 vps; p=3×10^−8^, two-sided Poisson exact test; Fig. 1B), with a consistent 1.2-fold enrichment in the case-control data (4,277 in 5,556 cases versus 6,149 in 8,809 controls; 0.80 vs. 0.68 vps; p=4×10^−7^, binomial exact test; Fig. 1B). Of note, in the *de novo* data, this top tier of missense variation shows stronger enrichment in cases than the middle tier of PTVs. The other two tiers of missense variation are not significantly enriched in cases (Table S3).

### Sex differences in ASD risk

The prevalence of ASD is higher in males than females. In line with previous observations of females with ASD carrying a higher genetic burden than males (De Rubeis et al., 2014), we observe a 2-fold enrichment of *de novo* PTVs in highly constrained genes in affected females (n=1,097) versus affected males (n=5,333) (p=3×10^−6^, two-sided Poisson exact test; Fig. 1B; Table S3). This result is consistent with the female protective effect (FPE) model, which postulates that females require an increased genetic load (in this case, high-liability PTVs) to reach the threshold for a diagnosis (Werling, 2016). The converse hypothesis is that risk variation has larger effects in males than in females so that females require a higher genetic burden to reach the same diagnostic threshold as males; however, across all classes of genetic variants, we observed no significant sex differences in trait liability, consistent with the FPE model (STAR Methods; Fig. 1C). In the absence of sex-specific differences in liability, we estimated the liability z-scores for different classes of variants across both sexes together (Fig. 1C; Table S3) and leveraged them to enhance gene discovery.

### ASD gene discovery

In previous risk gene discovery efforts, we used the Transmitted And *De novo* Association (TADA) model (He et al., 2013) to integrate protein-truncating and missense variants that are *de novo*, inherited, or from case-control populations and to stratify autosomal genes by FDR for association. Here, we update the TADA model to include pLI score as a continuous metric for PTVs, and MPC score as a two-tiered metric (≥2, 1-2) for missense variants (STAR Methods; Fig. S2; Fig. S3). From family data we include *de novo* PTVs as well as *de novo* missense variants, while for case-control we include only PTVs; we do not include inherited variants due to the limited liabilities observed (Fig. 1C). These modifications result in an enhanced TADA model with greater sensitivity and accuracy than the original model (Fig. 2A; STAR Methods).

**Figure 2.**
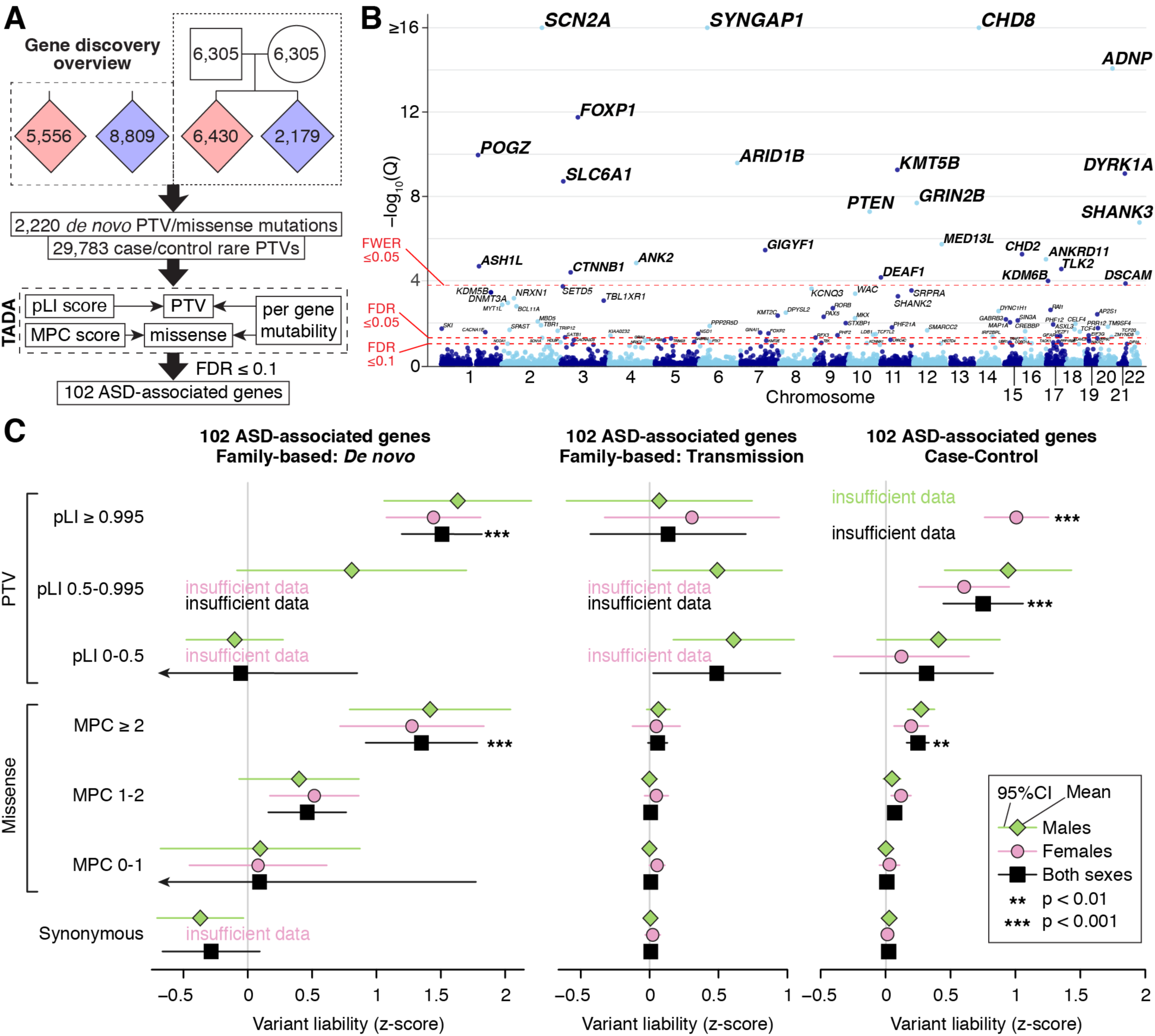
Gene discovery in the ASC cohort. **A**, WES data from 35,584 samples are entered into a Bayesian analysis framework (TADA) that incorporates pLI score for PTVs and MPC score for missense variants. **B**, The model identifies 102 autosomal genes associated with ASD at a false discovery rate (FDR) threshold of ≤ 0.1, which is shown on the y-axis of this Manhattan plot with each point representing a gene. Of these, 78 exceed the threshold of FDR ≤ 0.05 and 26 exceed the threshold family-wise error rate (FWER) ≤ 0.05. **C**, Repeating our ASD trait liability analysis (Fig. 1C) restricted to variants observed within the 102 ASD-associated genes only. Statistical tests: B, TADA; C, Binomial Exact Test (BET) for most contrasts; exceptions were “both” and “case-control”, for which Fisher’s method for combining BET p-values for each sex and, for case-control, each population, was used; p-values corrected for 168 tests are shown.

Our refined TADA model identifies 102 ASD risk genes at FDR ≤ 0.1, of which 78 meet the more stringent threshold of FDR ≤ 0.05, with 26 significant after Bonferroni correction (Fig. 2B; Table S4). By simulation experiments (Supplemental Methods), we demonstrate the reliable performance of our model, in particular showing that FDR is properly calibrated (Fig. S2). Of the 102 ASD-associated genes, 60 were not discovered by our earlier analyses (De Rubeis et al., 2014; Sanders et al., 2015), including 31 that have not been implicated in autosomal dominant neurodevelopmental disorders and were not significantly enriched for *de novo* and/or rare variants in previous studies, and that can therefore be considered novel (Table S5). The patterns of liability seen for these 102 genes are similar to that seen over all genes (compare Fig. 2C versus Fig. 1C), although the effects of variants are uniformly larger, as would be expected for this selected list of putative risk genes that would be enriched for true risk variants.

### Patterns of mutations in ASD genes

Within the set of observed mutations, the ratio of PTVs to missense mutations varies substantially between genes (Fig. 3A). Some genes, such as *ADNP*, reach our association threshold through PTVs alone, amongst which three genes have a significant excess of PTVs, relative to missense mutations, in the current dataset, based on gene mutability: *SYNGAP1*, *DYRK1A*, and *ARID1B* (p < 0.0005, binomial test). Because of the increased cohort size and availability of the MPC metric, we are also able for the first time to associate genes with ASD based primarily on *de novo* missense variation. We therefore examined four genes with four or more *de novo* missense variants (MPC ≥ 1) in ASD cases and one or no PTVs: *DEAF1*, *KCNQ3*, *SCN1A*, and *SLC6A1* (Fig. 3A; Table S6).

**Figure 3.**
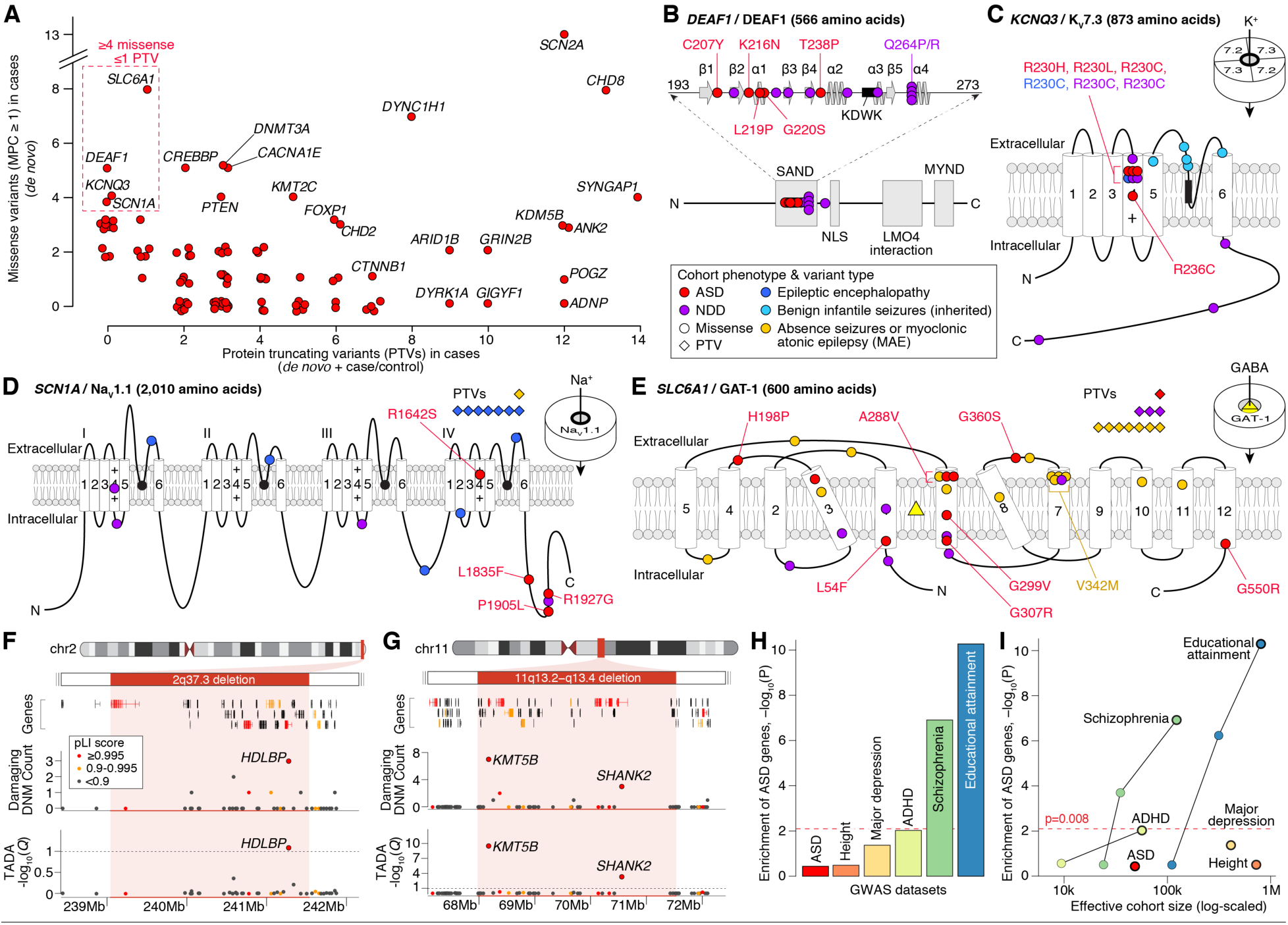
Genetic characterization of ASD genes. **A**, Count of PTVs versus missense variants (MPC ≥ 1) in cases for each ASD-associated gene (red points, selected genes labeled). These counts reflect the data used by TADA for association analysis: de novo and case/control data for PTVs; de novo only for missense. **B**, Location of ASD de novo missense variants in DEAF1. The five ASD mutations (marked in red) are in the SAND DNA-binding domain (amino acids 193-273, spirals show alpha helices, arrows show beta sheets, KDWK is the DNA-binding motif) alongside ten variants observed in NDD, several of which have been shown to reduce DNA binding, including Q264P and Q264R (Chen et al., 2017; Heyne et al., 2018; Vulto-van Silfhout et al., 2014). **C**, Location of ASD missense variants in KCNQ3. All four ASD variants were located in the voltage sensor (fourth of six transmembrane domains), with three in the same residue (R230), including the gain-of-function R230C mutation observed in NDD (Heyne et al., 2018). Five inherited variants observed in benign infantile seizures are shown in the pore loop (Landrum et al., 2014; Maljevic et al., 2016). **D**, Location of ASD missense variants in SCN1A, alongside 17 de novo variants in NDD and epilepsy (Heyne et al., 2018). **E**, Location of ASD missense variants in SLC6A1, alongside 31 de novo variants in NDD and epilepsy (Heyne et al., 2018; Johannesen et al., 2018). **F**, Subtelomeric 2q37 deletions are associated with facial dysmorphisms, brachydactyly, high BMI, neurodevelopmental delay, and ASD (Leroy et al., 2013). While three genes within the locus have a pLI score ≥ 0.995, only HDLBP is associated with ASD. **G**, Deletions at the 11q13.2−q13.4 locus have been observed in NDD, ASD, and otodental dysplasia (Coe et al., 2014; Cooper et al., 2011). Five genes within the locus have a pLI score ≥ 0.995, including two ASD genes: KMT5B and SHANK2. **H**, Assessment of gene-based enrichment, via MAGMA, of 102 ASD genes against genome-wide significant common variants from six GWAS. **I**, Gene-based enrichment of 102 ASD genes in multiple GWAS as a function of effective cohort size. The GWAS used for each disorder in ‘I’ has a black outline. Statistical tests: F, G, TADA; H, I, MAGMA.

We observe five *de novo* missense variants and no PTVs in *DEAF1*, which encodes a self-dimerizing transcription factor involved in neuronal differentiation (Bottomley et al., 2001). All five missense variants reside in the SAND domain (Fig. 3B), which is critical for both dimerization and DNA binding (Bottomley et al., 2001; Jensik et al., 2004). A similar pattern of SAND domain missense enrichment is observed in individuals with intellectual disability, speech delay, and behavioral abnormalities (Chen et al., 2017; Heyne et al., 2018; Vulto-van Silfhout et al., 2014).

Four *de novo* missense variants and no PTVs are observed in *KCNQ3*, which encodes a subunit of a neuronal voltage-gated potassium channel (Fig. 3C). All four variants modify arginine residues in the voltage-sensing fourth transmembrane domain, with three at a single residue previously characterized as gain-of-function in NDD (R230C, Fig. 3C) (Miceli et al., 2015). These data suggest gain-of-function mutations in *KCNQ3* also confer risk to ASD.

*SCN1A* encodes a voltage-gated sodium channel and has been associated, predominantly through PTVs, with Dravet syndrome (Claes et al., 2001), a form of progressive epileptic encephalopathy which often meets diagnostic criteria for ASD (Rosander and Hallbook, 2015). We observe four *de novo* missense variants and no PTVs in *SCN1A* (Fig. 3A; Table S6), with three located in the C-terminus (Fig. 3D), and all four cases are reported to have seizures.

The gene *SLC6A1* encodes a voltage-gated GABA transporter and has been associated with developmental delay and cognitive impairment (Deciphering Developmental Disorders, 2017; Heyne et al., 2018), as well as myoclonic atonic epilepsy and absence seizures (Johannesen et al., 2018). Here, we extend the phenotypic spectrum to include ASD, through the observation of eight *de novo* missense variants and one PTV, all in cases (Fig. 3E). Four of these variants reside in the sixth transmembrane domain, with one recurring in two independent cases (A288V). Five of the six cases with available information on history of seizure had seizures, and all four cases with available data on cognitive performance have intellectual disability.

### ASD genes within recurrent copy number variants (CNVs)

Large CNVs represent another important source of risk for ASD (Sebat et al., 2007), but these genomic disorder (GD) segments can include dozens of genes, complicating the identification of driver gene(s) within these regions. We sought to determine whether the 102 ASD genes could nominate driver genes within GD regions. We first curated a consensus GD list from nine sources, totaling 823 protein-coding genes in 51 autosomal GD loci associated with ASD or ASD-related phenotypes, including NDD (Table S7).

Within the 51 GDs, 12 GD loci encompassed 13 ASD-associated genes (Table S7), which is greater than expected by chance when simultaneously controlling for number of genes, PTV mutation rate, and brain expression levels per gene (2.3-fold increase; p=2.3×10^−3^, permutation). These 12 GD loci divided into three groups: 1) the overlapping ASD gene matched the consensus driver gene, e.g., *SHANK3* for Phelan-McDermid syndrome (Soorya et al., 2013); 2) an ASD gene emerged that did not match the previously predicted driver gene(s) within the region, such as *HDLBP* at 2q37.3 (Fig. 3F), where *HDAC4* has been hypothesized as a driver gene in some a analyses (Williams et al., 2010); and 3) no previous driver gene had been established within the GD locus, such as *BCL11A* at 2p15-p16.1. One GD locus, 11q13.2-q13.4, had two of our 102 genes (*SHANK2* and *KMT5B*, Fig. 3G), highlighting that GDs can result from risk conferred by multiple genes, potentially including genes with small effect sizes that we are underpowered to detect.

### Relationship of ASD genes with GWAS signal

Common variation plays an important role in ASD risk, and recent genome-wide association studies (GWAS) have revealed a handful of ASD-associated loci (Grove et al., 2019). Thus, we asked if common genetic variation within or near the 102 identified genes (within 10 Kb) influences ASD risk or other traits related to ASD risk. We note that, among the first five genome-wide significant ASD hits from the current largest GWAS (Grove et al., 2019), *KMT2E* is a “double hit”—implicated by both the GWAS and the list of 102 FDR ≤ 0.1 genes described here.

We therefore ran a gene set enrichment analysis of our 102 ASD-associated genes against GWAS summary statistics using MAGMA (de Leeuw et al., 2015) to integrate the signal for those statistics over each gene using brain-expressed protein-coding genes as our background. We used results from six GWAS datasets: ASD, schizophrenia (SCZ), major depressive disorder (MDD), and attention deficit hyperactivity disorder (ADHD), which are all positively genetically correlated with ASD and with each other; educational attainment (EA), which is positively correlated with ASD and negatively correlated with schizophrenia and ADHD; and human height as a negative control (Table S8) (Demontis et al., 2018; Grove et al., 2019; Lee et al., 2018; Neale et al., 2010; Okbay et al., 2016; Rietveld et al., 2013; Ripke et al., 2013a; Ripke et al., 2011; Ripke et al., 2013b; Schizophrenia Working Group of the Psychiatric Genomics, 2014; Wray et al., 2018; Yengo et al., 2018; Zheng et al., 2017). Correcting for six analyses, we observed significant enrichment only for SCZ and EA (Fig. 3H). Curiously, the ASD and ADHD GWAS signals were not enriched in the 102 ASD genes. Although in some ways these results are counterintuitive, one obvious confounder is power (Fig. 3I). Effective cohort sizes for the SCZ, EA, and height GWAS dwarf that for ASD, and the quality of GWAS signal strongly increases with sample size. Thus, for results from well-powered GWAS, it is reassuring that there is no signal for height, yet clearly detectable signal for two traits genetically correlated to ASD: SCZ and EA.

### Relationship between ASD and other neurodevelopmental disorders

Family studies yield high heritability estimates in ASD (Yip et al., 2018), but comparable estimates of heritability in severe NDD are lower (Reichenberg et al., 2016). Consistent with these observations, exome studies identify a higher frequency of disruptive *de novo* variants in severe NDD than in ASD (Deciphering Developmental Disorders, 2017). Because of the 30% co-morbidity between ASD subjects and intellectual disability/NDD, it is unsurprising that many genes are associated with both disorders (Pinto et al., 2010). Distinguishing genes that, when disrupted, lead to ASD more frequently than NDD may shed new light on how atypical neurodevelopment maps onto the core deficits of ASD.

To partition the 102 ASD genes in this manner, we compiled data from 5,264 trios ascertained for severe NDD (Table S9) and compared the relative frequency, *R*, of disruptive *de novo* variants (which we define as PTVs or missense variants with MPC ≥ 1) in ASD- or NDD-ascertained trios. Genes with *R* > 1 were classified as ASD-predominant (ASD_P_, 50 genes), while those with R < 1 were classified as ASD with NDD (ASD_NDD_, 49 genes). An additional three genes were assigned to the ASD_P_ group on the basis of case-control data, totaling 53 ASD_P_ genes (Fig. 4A). For this partition, we then evaluated transmission of rare PTVs (relative frequency < 0.001) from parents to their affected offspring: for ASD_P_ genes, 44 such PTVs were transmitted and 18 were not (p=0.001, transmission disequilibrium test [TDT]), whereas, for ASD_NDD_ genes, 14 were transmitted and 8 were not (p=0.29; TDT). The frequency of PTVs in parents is significantly greater in ASD_P_ genes (1.17 per gene) than in ASD_NDD_ genes (0.45 per gene; p=6.6×10^−6^, binomial test), while the frequency of *de novo* PTVs in probands is not markedly different between the two groups (95 in ASD_P_ genes, 121 in ASD_NDD_ genes, p=0.07, binomial test with probability of success = 0.503 [PTV in ASD_P_ gene]). The paucity of inherited PTVs in ASD_NDD_ genes is consistent with greater selective pressure acting against disruptive variants in these genes.

**Figure 4.**
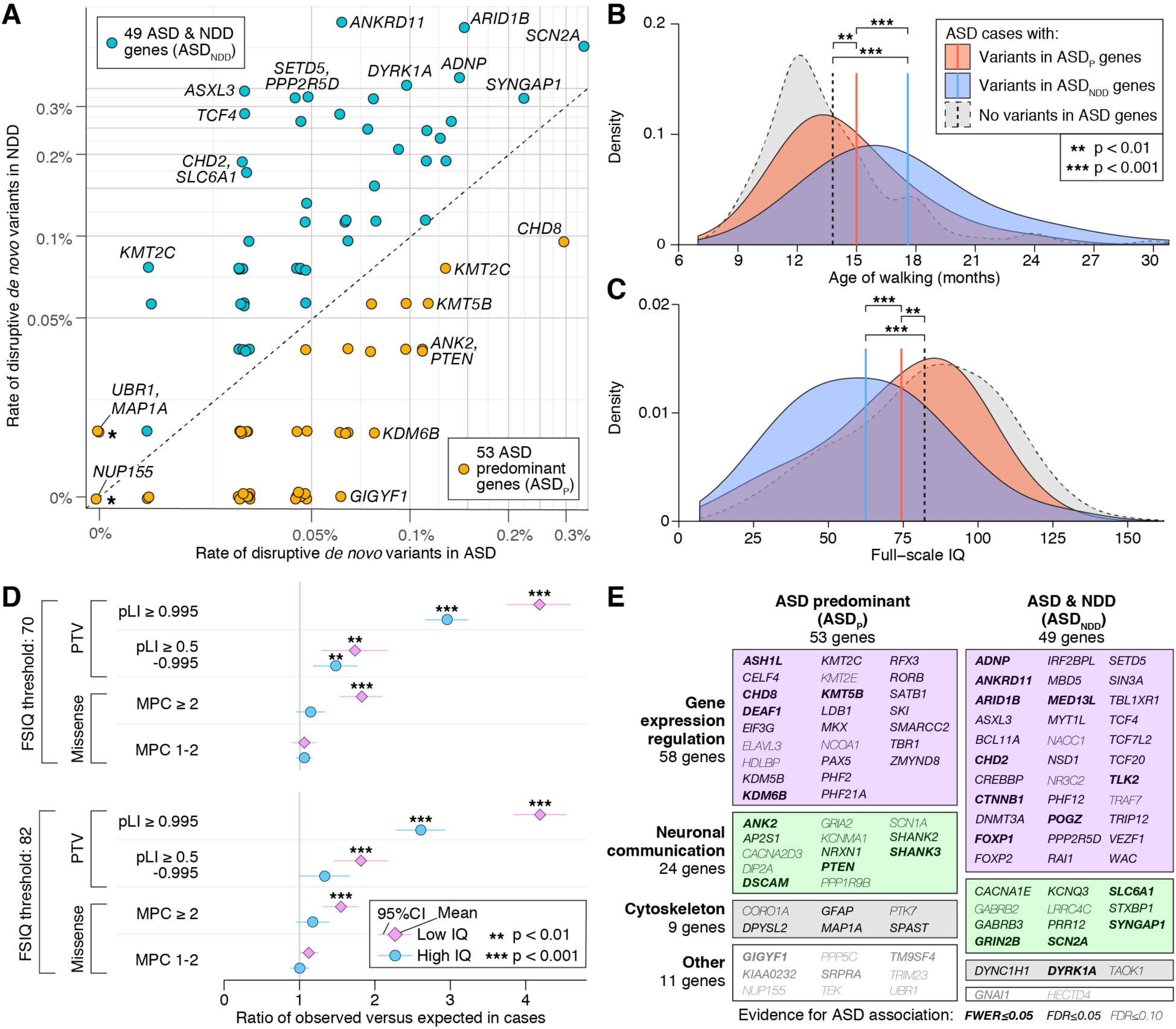
Phenotypic and functional categories of ASD-associated genes. **A**, The frequency of disruptive de novo variants (e.g. PTVs or missense variants with MPC ≥ 1) in ASD-ascertained and NDD-ascertained cohorts (Table S4) is shown for the 102 ASD-associated genes (selected genes labeled). Fifty genes with a higher frequency in ASD are designated ASD-predominant (ASD_P_), while the 49 genes more frequently mutated in NDD are designated as ASD_NDD_. Three genes marked with a star (UBR1, MAP1A, and NUP155) are included in the ASD_P_ category on the basis of case-control data (Table S4), which are not shown in this plot. **B**, ASD cases with disruptive de novo variants in ASD genes show delayed walking compared to ASD cases without such de novo variants, and the effect is greater for those with disruptive de novo variants in ASD_NDD_ genes. **C**, Similarly, cases with disruptive de novo variants in ASD_NDD_ genes and, to a lesser extent, ASD_P_ genes have a lower full-scale IQ than other ASD cases. **D**, Despite the association between de novo variants in ASD genes and cognitive impairment shown in ‘C’, an excess of disruptive de novo variants is observed in cases without intellectual disability (FSIQ ≥ 70) or with an IQ above the cohort mean (FSIQ ≥ 82). **E**, Along with the phenotypic division (A), genes can also be classified functionally into four groups (gene expression regulation (GER), neuronal communication (NC), cytoskeleton, and other) based on gene ontology and research literature. The 102 ASD risk genes are shown in a mosaic plot divided by gene function and, from ‘A’, the ASD vs. NDD variant frequency, with the area of each box proportional to the number of genes. Statistical tests: B, C, t-test; D, chi-square with 1 degree of freedom.

Consistent with this partition, ASD subjects who carry disruptive *de novo* variants in ASD_NDD_ genes walk 2.6 ± 1.2 months later (Fig. 4B; p=2.3×10^−5^, t-test, df=251) and have an IQ 11.9 ± 6.0 points lower (Fig. 4C; p=1.1×10^−4^, two-sided t-test, df=278), on average, than ASD subjects with disruptive *de novo* variants in ASD_P_ genes (Table S10). Both sets of subjects differ significantly from the rest of the cohort with respect to IQ and age of walking (Fig. 4B, 4C; Fig. S4; Table S10). Thus, the data support some overall distinction between the genes identified in ASD and NDD *en masse*, although our current analyses are not powered for variant-level or gene-level resolution.

### Burden of mutations in ASD as a function of IQ

Of the 6,430 ASD probands, 3,010 had a detected *de novo* variant and either a recorded full-scale IQ or a clinical assessment of ID. We partitioned these subjects into those with IQ ≥ 70 (69.4%) versus those with IQ < 70 (30.6%), then characterized the burden of *de novo* variants within these groups. ASD subjects in the lower IQ group carry a greater burden of *de novo* variants, relative to both expectation and the high IQ group, in the two top tiers of PTVs and the top tier of missense variants (Fig. 4D). Excess burden relative to expectation is also observed in the two top PTV tiers for the high IQ group (Fig. 4D). Similar patterns were observed partitioning the sample at IQ ≥ 82 (53.7%) versus IQ < 82 (46.3%), which was the mean IQ after removing subjects who carry disruptive variants in the 102 ASD genes (Fig. 4C). Finally, we observe excess burden in the high IQ group when considering the 102 ASD genes only, as documented by model-driven simulations accounting for selection bias due to an FDR threshold (STAR Methods). Thus, excess burden is not limited to low IQ cases, supporting the idea that *de novo* variants do not solely impair cognition (Robinson et al., 2014).

### Functional dissection of ASD genes

Past WES analyses have identified two major functional groups of ASD genes: those involved in gene expression regulation (GER), including chromatin regulators and transcription factors, and those involved in neuronal communication (NC), including synaptic function (De Rubeis et al., 2014). A simple gene ontology enrichment analysis with the new list of 102 ASD genes replicates this finding, identifying 16 genes in category GO:0006357 “regulation of transcription from RNA polymerase II promoter” (5.7-fold enrichment, FDR=6.2×10^−6^) and 9 genes in category GO:0007268: “synaptic transmission” (5.0-fold enrichment, FDR=3.8×10^−3^). We used a combination of gene ontology and primary literature research to assign additional genes to the GER (58 genes) and NC (24 genes) categories for further analyses (STAR Methods; Table S11; Fig. 4E). We also see the emergence of a new functional group of nine genes implicated in category GO:0007010 “cytoskeleton organization”. The remaining 11 genes are described as “Other” (Table S11 and Fig. 4E), many of which have roles in signaling cascades and/or ubiquitination.

### ASD genes are expressed early in brain development

The 102 ASD genes can thus be subdivided by functional role (58 GER genes, 24 NC genes) and phenotypic impact (53 ASD_P_ genes, 49 ASD_NDD_ genes) to give five gene sets (including the set of all 102). We first evaluated enrichment of these five gene sets in the 53 tissues with bulk RNA-seq data in the Genotype-Tissue Expression (GTEx) resource (GTEx-Consortium, 2017). To enhance tissue-specific resolution, we selected genes that were expressed in one tissue at a significantly higher level than the remaining 52 tissues, specifically log fold-change > 0.5 and FDR < 0.05 (*t*-test). Subsequently, we assessed over-representation of each ASD gene set within 53 sets of genes expressed in each tissue relative to a background of all tissue-specific genes in GTEx. At a multiple-testing threshold of p ≤ 9×10^−4^, reflecting 53 tissues, enrichment was observed in 11 of the 13 brain regions, with the strongest enrichment in cortex (∩=30 genes; p=3×10^−6^; OR=3.7; Fig. 5A) and cerebellar hemisphere (∩=48 genes; p=3×10^−6^; OR=2.9; Fig. 5A). Of the four gene subsets, NC genes were the most highly enriched in cortex (∩=17/23 genes; p=3×10^−11^; OR=25; Fig. 5A), while GER genes were the least enriched (∩=10/58 genes; p=0.36; OR=1.8; Fig. 5A).

**Figure 5.**
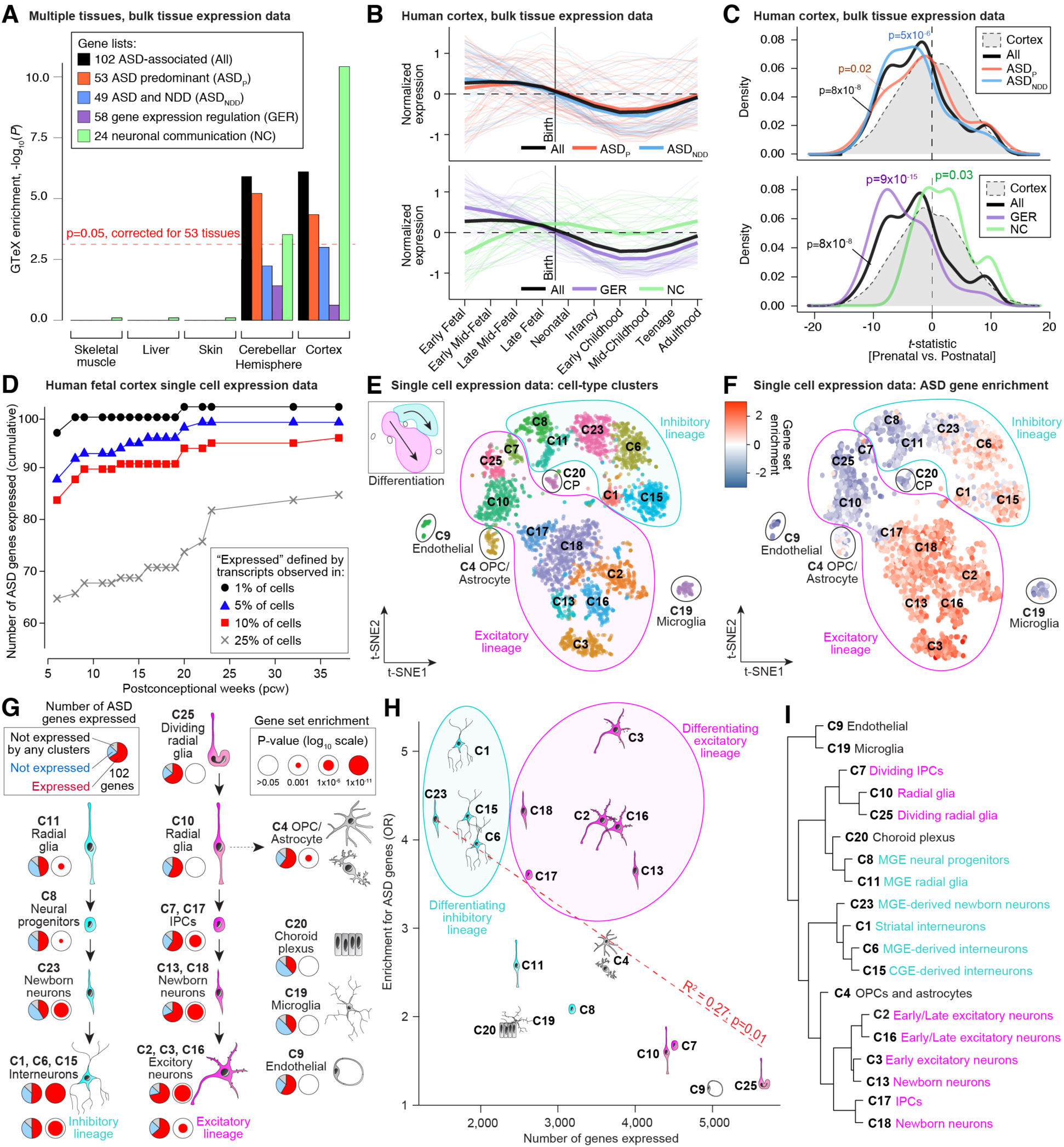
Analysis of 102 ASD-associated genes in the context of gene expression data. **A**, GTEx bulk RNA-seq data from 53 tissues was processed to identify genes enriched in specific tissues. Gene set enrichment was performed for the 102 ASD genes and four subsets (ASD_P_, ASD_NDD_, GER, NC) for each tissue. Five representative tissues are shown here, including cortex, which has the greatest degree of enrichment (OR=3.7; p=2.6×10^−6^). **B**, BrainSpan bulk RNA-seq data across 10 developmental stages was used to plot the normalized expression of the 101 brain-expressed ASD genes across development, split by the four subsets. **C**, A t-statistic was calculated comparing prenatal to postnatal expression in the BrainSpan data. The t-statistic distribution of 101 ASD-associated genes (excluding PAX5, which is not expressed in the cortex) shows a prenatal bias (p=8×10^−8^) for GER genes (p=9×10^−15^), while NC genes are postnatally biased (p=0.03). **D**, The cumulative number of ASD-associated genes expressed in RNA-seq data for 4,261 cells collected from human forebrain across prenatal development. **E**, t-SNE analysis identifies 19 clusters with unambiguous cell type in this single-cell expression data. **F**, The enrichment of the 102 ASD-associated genes within cells of each type is represented by color. The most consistent enrichment is observed in maturing and mature excitatory (bottom center) and inhibitory (top right) neurons. **G**, The developmental relationships of the 19 clusters are indicated by black arrows, with the inhibitory lineage shown on the left (cyan), excitatory lineage in the middle (magenta), and non-neuronal cell types on the right (grey). The proportion of the 102 ASD-associated genes observed in at least 25% of cells within the cluster is shown by the pie chart, while the log-transformed Bonferroni corrected p-value of gene set enrichment is shown by the size of the red circle. **H**, The relationship between the number of cells in the cluster (x-axis) and the p-value for ASD gene enrichment (y-axis) is shown for the 19 cell type clusters. Linear regression indicates that clusters with few expressed genes (e.g. C23 newborn inhibitory neurons) have higher p-values than clusters with many genes (e.g. C25 radial glia). **I**, The relationship between the 19 cell type clusters using hierarchical clustering based on the 10% of genes with the greatest variability among cell types. Statistical tests: A, t-test; C, Wilcoxon test; E, F, H, I, Fisher’s Exact Test.

We next leveraged BrainSpan human neocortex bulk RNA-seq data (Li et al., 2018) to assess enrichment of ASD genes across development (Fig. 5B, 5C). Of the 17,484 autosomal protein-coding genes assessed for association, 13,735 (78.5%) were expressed in the cortex (RPKM ≥ 0.5 in 80% of samples of at least one cortical region and developmental period). Of the 102 ASD genes, only the cerebellar transcription factor *PAX5* (FDR=0.005, TADA) was not expressed in the cortex (78 expected; p=1×10^−9^, binomial test). Compared to other genes expressed in the cortex, the remaining 101 ASD genes are expressed at higher levels during prenatal development (Fig. 5B). To quantify this pattern, we developed a *t*-statistic that assesses the relative prenatal vs. postnatal expression of each of the 13,735 expressed genes. Using this metric, the 101 cortically-expressed ASD genes showed enrichment in the prenatal cortex (p=8×10^−8^, Wilcoxon test; Fig. 5C). The ASD_P_ and ASD_NDD_ gene sets showed similar patterns (Fig. 5B), though the prenatal bias *t*-statistic was slightly more pronounced for the ASD_NDD_ group (p=5×10^−6^, Wilcoxon test; Fig. 5C). The GER genes reach their highest levels during early to late fetal development (Fig. 5B) with a marked prenatal bias (p=9×10^−15^, Wilcoxon test; Fig. 5C), while the NC genes are highest between late midfetal development and infancy (Fig. 5B) and show a trend towards postnatal bias (p=0.03, Wilcoxon test; Fig. 5C). Thus, in keeping with prior analyses (Chang et al., 2014; Parikshak et al., 2013; Willsey et al., 2013; Xu et al., 2014), the ASD genes are expressed at high levels in human cortex and are expressed early in brain development. The differing expression patterns of GER and NC genes may reflect two distinct periods of ASD susceptibility during development or a single susceptibility period when both functional gene sets are highly expressed in mid- to late fetal development.

### ASD genes are enriched in maturing inhibitory and excitatory neurons

Prior analyses have implicated excitatory glutamatergic neurons in the cortex and medium spiny neurons in the striatum in ASD (Chang et al., 2014; Parikshak et al., 2013; Willsey et al., 2013; Xu et al., 2014). Here, we perform a more direct assessment, examining expression of the 102 ASD-associated genes in an existing single-cell RNA-seq dataset of 4,261 cells from the prenatal human forebrain (Nowakowski et al., 2017), ranging from 6 to 37 post-conception weeks (pcw) with an average of 16.3 pcw (Table S12).

Following the logic that only genes that were expressed could mediate ASD risk when disrupted, we divided the 4,261 cells into 17 bins by developmental stage and assessed the cumulative distribution of expressed genes by developmental endpoint (Fig. 5D). For each endpoint, a gene was defined as expressed if at least one transcript mapped to this gene in 25% or more of cells for at least one pcw stage. By definition, more genes were expressed as fetal development progressed, with 4,481 genes expressed by 13 pcw and 7,171 genes expressed by 37 pcw. While the majority of ASD genes were expressed at the earliest developmental stages (e.g. 68 of 102 at 13 pcw), the most dramatic increase in the number of genes expressed occurred during midfetal development (70 by 19 pcw, rising to 81 by 23 pcw), consistent with the BrainSpan bulk-tissue data (Fig. 5B, 5C). More liberal thresholds for gene expression resulted in higher numbers of ASD genes expressed (Fig. 5D), but the patterns of expression were similar across definitions and when considering gene function or cell type (Fig. S5).

To investigate the cell types implicated in ASD, we considered 25 cell type clusters identified by t-distributed stochastic neighbor embedding (t-SNE) analysis, of which 19 clusters, containing 3,839 cells, were unambiguously associated with a cell type (Nowakowski et al., 2017) (Fig. 5E, Table S12), and were used for enrichment analysis. Within each cell type cluster, a gene was considered expressed if at least one of its transcripts was detected in 25% or more cells; 7,867 protein coding genes met this criterion in at least one cluster. By contrasting one cell type to the others, we observed enrichment for the 102 ASD genes in maturing and mature neurons of the excitatory and inhibitory lineages (Fig. 5F, 5G) but not in non-neuronal lineages. Early excitatory neurons (C3) expressed the most ASD genes (∩=72 genes, OR=5.0, p < 1×10^−10^, Fisher’s exact test [FET]), while choroid plexus (C20) and microglia (C19) expressed the fewest ASD genes (∩=39 genes, p=0.09 and 0.137, respectively, FET); 14 genes were not expressed in any cluster (Fig. 5G). Within the major neuronal lineages, early excitatory neurons (C3) and striatal interneurons (C1) showed the greatest degree of gene set enrichment (∩=72 and ∩=51 genes, p < 1×10^−10^, FET; Fig. 5F, 5G; Table S12). Overall, maturing and mature neurons in the excitatory and inhibitory lineages showed a similar degree of enrichment, while those in the excitatory lineage expressed the most ASD genes, paralleling the larger numbers of genes expressed in excitatory lineage cells (Fig. 5H). The only non-neuronal cell type with significant enrichment for ASD genes was oligodendrocyte progenitor cells (OPCs) and astrocytes (C4; ∩=62 genes, OR=2.8, p=8×10^−5^, FET). Of the 60 ASD genes expressed in OPCs, 58 overlapped with radial glia, which may reflect shared developmental origins rather than an independent enrichment signal. In contrast to post-mortem findings in adult ASD brains (Gandal et al., 2018; Voineagu et al., 2011), no enrichment was observed in microglia. To validate the t-SNE clusters, we selected 10% of the expressed genes showing the greatest variability among the cell types and performed hierarchical clustering (Fig. 5I). This recaptured the division of these clusters by lineage (excitatory vs. inhibitory) and by development stage (radial glia and progenitors vs. neurons).

### Functional relationships among ASD genes and prediction of novel risk genes

The ASD genes show convergent functional roles (Fig. 4E) and expression patterns in the cortex (Fig. 5B). It is therefore reasonable to hypothesize that genes co-expressed with these ASD genes might have convergent or auxiliary function and thus contribute to risk. We have previously developed the Discovering Association With Networks (DAWN) approach to integrate TADA scores of genetic association and gene co-expression data in order to uncover more risk genes. By using the TADA results and BrainSpan gene co-expression data from the midfetal human cortex, DAWN yields 138 genes (FDR ≤ 0.005), including 83 genes that are not captured by TADA, with 69 of these 83 correlated with many other genes (Fig. 6A; Table S13). Of the 83, 19 are implicated in neurodevelopmental disorders, and seven of these have autosomal recessive inheritance (Table S5, S13). Of the 138 genes, 38 are expressed in excitatory cell types (p < 1.6×10^−4^, FET), 25 are also expressed in inhibitory cell types (p < 7.9×10^−4^, FET), and many play a role in GER or NC (Fig. 6A).

**Figure 6.**
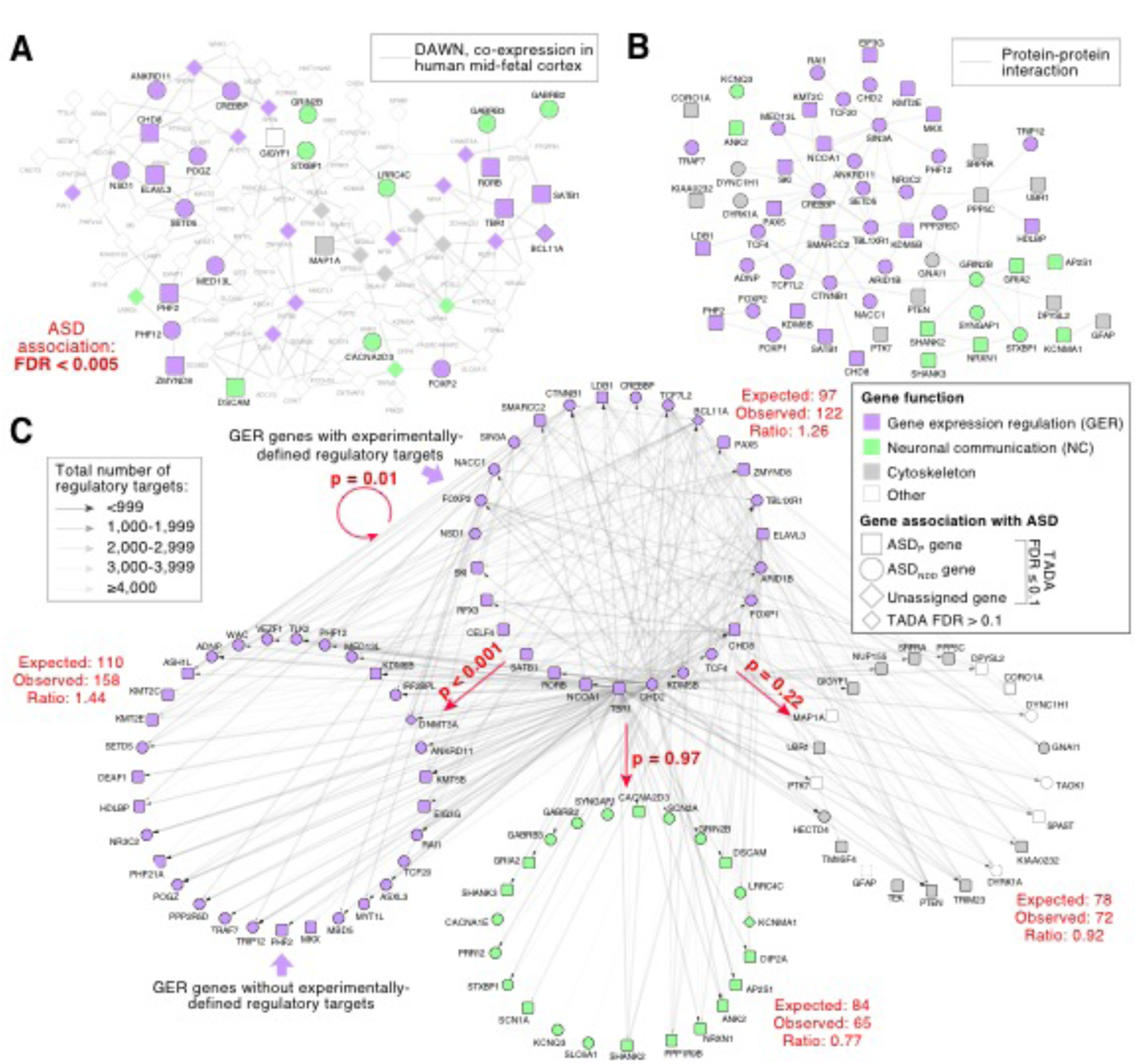
Functional relationships of ASD risk genes. **A**, ASD association data from TADA (Table S4) is integrated with co-expression data from the midfetal human brain to implicate additional genes in ASD using DAWN (Discovering Association With Networks). The top 138 genes that share edges are shown (FDR ≤ 0.005). **B**, ASD-associated genes form a single protein-protein interaction network with more edges than expected by chance (p=0.01). **C**, Experimental data, obtained using ChIP and CLIP methods across multiple species and a wide range of neuronal and non-neuronal tissues types, identified the regulatory targets of 26 GER genes (top circle). These data were used to assess whether three functionally-defined groups of ASD-associated genes were enriched for regulatory targets, represented as arrows, weighted by the total number of regulatory targets for the GER gene. The expected number of targets in each functional group was estimated by permutation, controlling for brain expression, de novo PTV mutation rate, and pLI. Statistical tests: A, DAWN; B, Permutation; C, Permutation.

To interpret gene co-expression and enrichment across a broader range of early developmental samples, we used Weighted Gene Coexpression Network Analysis (WGCNA) to assess spatiotemporal co-expression from 177 high-quality BrainSpan samples aged 8 pcw to 1 year. WGCNA yielded 27 early developmental co-expression modules, two of which show significant over-representation of 102 ASD genes after correction for multiple testing (Fig. S6, Table S14): M4 for the NC gene set (∩=5 genes, OR=13.7, p=0.002, FET); and M25 for all 102 ASD genes (∩=17 genes, OR=12.1, p=3×10^−11^, FET), although driven by the GER gene set (∩=17 genes, OR=26.2, p=9×10^−16^, FET). With regard to single-cell gene expression, genes in the NC-specific M4 showed greatest enrichment in maturing neurons, both excitatory and inhibitory (p < 0.001 for each of 6 neuronal cell types, FET), whereas genes in M25 showed enrichment across all 19 cell types (p < 0.001 for all cell types, FET). The DAWN associated genes are enriched in M3 (∩=10, OR=10.1, p=5×10^−6^, FET) and M25 (∩=7, OR=5.9, p=0.004, FET) but not M4, although expression of genes in M4 are highly correlated with those of M3 during early development, and both are highly expressed prenatally. Comparing our gene modules to previously published candidate ASD gene networks obtained using WGCNA (Parikshak et al., 2013) shows that our M4 strongly overlaps with previously identified M16 (p=5.5×10^−69^, FET), and our M24 overlaps with previously identified M2 (p=1.3×10^−239^, FET) (note, however that both studies make use of BrainSpan data so the overlap is not unexpected but helps to relate the results from the two studies).

To explore whether GER and NC gene sets interact more than would be expected by chance, we analyzed protein-protein interaction (PPI) networks (Fig. 6B; Table S15) and found they do not: there was a significant excess of interactions among all ASD genes (∩=82 genes, *p*=0.02, FET), GER genes (∩=49 genes, *p*=0.006, FET), and NC genes (∩=12 genes, *p*=0.03, FET), but not among GER and NC genes (∩=2 genes, *p*=1.00, FET). Nor do GER genes regulate the NC genes, according to our analyses, although GER-GER regulation was enriched (Table S16, Fig. 6C). Even *CHD8*, the most prominent and well characterized ASD GER gene, did not regulate NC genes more than expected by chance (Fig. S7).

## Discussion

By characterizing rare *de novo* and inherited coding variation from 35,584 individuals, including 11,986 ASD cases, we implicate 102 genes in risk for ASD at FDR ≤ 0.1 (Fig. 2), of which 31 are novel risk genes. Notably, analyses of this set of risk genes lead to novel genetic, phenotypic, and functional findings. Evidence for several of the genes is driven by missense variants, including confirmed gain-of-function mutations in the potassium channel *KCNQ3* and patterns that may similarly reflect gain-of-function or altered function in *DEAF1*, *SCN1A*, and *SLC6A1* (Fig. 3). Further, we strengthen evidence for driver genes in genomic disorder loci and we propose a new driver gene (*BCL11A*) for the recurrent CNV at 2p15-p16.1. By evaluating GWAS results for ASD and related phenotypes and asking whether their common variant association signals overlap significantly with the 102 risk genes, we find substantial enrichment of GWAS signal for two traits genetically correlated with ASD—schizophrenia and educational attainment. For ASD itself, however, this enrichment is not significant, likely due to the limited power of the ASD GWAS. Despite this, *KMT2E* is significantly associated with ASD by both common and rare risk variation.

We perform a genetic partition between genes predominantly conferring liability for ASD (ASD_P_) and genes imparting risk to both ASD and NDD (ASD_NDD_). Two lines of evidence support the partition: first, cognitive impairment and motor delay are more frequently observed in our subjects—all ascertained for ASD—with mutations in ASD_NDD_ than in ASD_P_ genes (Fig. 4B, 4C); second, we find that inherited variation plays a lesser role in ASD_NDD_ than in ASD_P_ genes. Together, these observations indicate that ASD-associated genes are distributed across a spectrum of phenotypes and selective pressure. At one extreme, gene haploinsufficiency leads to global developmental delay, with impaired cognitive, social, and gross motor skills leading to strong negative selection (e.g. *ANKRD11*, *ARID1B*). At the other extreme, gene haploinsufficiency leads to ASD, but there is a more modest involvement of other developmental phenotypes and selective pressure (e.g. *GIGYF1*, *ANK2*). This distinction has important ramifications for clinicians, geneticists, and neuroscientists, because it suggests that clearly delineating the impact of these genes across neurodevelopmental dimensions may offer a route to deconvolve the social dysfunction and repetitive behaviors that define ASD from more general neurodevelopmental impairment. Larger cohorts will be required to reliably identify specific genes as being enriched in ASD compared to NDD.

Single-cell gene expression data from the developing human cortex implicate mid-to-late fetal development and maturing and mature neurons in both excitatory and inhibitory lineages (Fig. 5). Expression of GER genes shows a prenatal bias, while expression of NC genes does not. Placing these results in the context of multiple non-exclusive hypotheses around the origins of ASD, it is intriguing to speculate that the NC ASD genes provide compelling support for E/I imbalance in ASD (Rubenstein and Merzenich, 2003) through direct impact on neurotransmission. However, as there was no support for a regulatory role for GER ASD genes on either NC or cytoskeletal ASD genes, additional mechanisms, having to do with cell migration and neurodevelopment, also appear to be at play. This might suggest that GER ASD genes impact E/I balance by altering the numbers of excitatory and inhibitory neurons in given regions of the brain. ASD must arise by phenotypic convergence amongst these diverse neurobiological trajectories, and further dissecting the nature of this convergence, especially in the genes that we have identified herein, is likely to hold the key to understanding the neurobiology that underlies the ASD phenotype.

## Acknowledgements

We thank the families who participated in this research, without whose contributions genetic studies would be impossible. This study was supported by the NIMH (U01s: MH100209 (to B.D.), MH100229 (to M.J.D.), MH100233 (to J.D.B), & MH100239 (to M.W.S.); U01s: MH111658 (to B.D.), MH111660 (to M.J.D.), MH111661 (to J.D.B), & MH111662 (to S.J.S. and M.W.S.); Supplement to U01 MH100233 (MH100233-03S1 to J.D.B.); R37 MH057881 (to B.D.); R01 MH109901 (to S.J.S. M.W.S., A.J.W.); R01 MH109900 (to K.R.); and, R01 MH110928 (to S.J.S., M.W.S., A.J.W.)), NHGRI (HG008895), Seaver Foundation, Simons Foundation (SF402281 to S.J.S., M.W.S., B.D., K.R.), and Autism Science Foundation (to S.J.S., S.L.B., E.B.R.). M.E.T. is supported by R01 MH115957 and Simons Foundation (SF573206) and R.C. is supported by NHGRI T32 HG002295-14 and NSF GRFP 2017240332. S.D.R. is supported by the Seaver Foundation. The iPSYCH project is funded by the Lundbeck Foundation (grant numbers R102-A9118 and R155-2014-1724) and the universities and university hospitals of Aarhus and Copenhagen. The Danish National Biobank resource at the Statens Serum Institut was supported by the Novo Nordisk Foundation. Sequencing of iPSYCH samples was supported by grants from the Simons Foundation (SFARI 311789 to M.J.D) and the Stanley Foundation. Other support for this study was received from the NIMH (5U01MH094432-02 to M.J.D). Computational resources for handling and statistical analysis of iPSYCH data on the GenomeDK and Computerome HPC facilities were provided by, respectively, Centre for Integrative Sequencing, iSEQ, Aarhus University, Denmark (grant to A.D.B), and iPSYCH. The Norwegian Mother and Child Cohort Study is supported by the Norwegian Ministry of Health and Care Services and the Ministry of Education and Research, NIH/NINDS (grant no.1 UO1 NS 047537-01 and grant no.2 UO1 NS 047537-06A1). We are grateful to all the participating families in Norway who take part in this ongoing cohort study and the Autism Birth Cohort Study. This work was also supported by the Research Council Norway grant number 185476 and Wellcome Trust grant number (098051]). For the collection of the cohort in Turin, the Italian Ministry for Education, University and Research (Ministero dell’Istruzione, dell’Università e della Ricerca - MIUR) funded the Department of Medical Sciences under the program “Dipartimenti di Eccellenza 2018 – 2022” Project code D15D18000410001. We also thank the Associazione “Enrico e Ilaria sono con noi” ONLUS and the Fondazione FORMA. The collection of the cohort in Santiago de Compostela, Spain (Angel Carracedo) was supported by the Fundación María José Jove. The collection of the cohort in Madrid, Spain (Mara Parellada) was funded by Instituto de Salud Carlos III (C.0001481, PI14/02103, PI17/00819) and IiSGM. The collection in Japan (Branko Aleksic) was supported by AMED under grant No. JP18dm0107087 and No. JP18dm0207005. The collection in Hong Kong (Brian H.Y. Chung) was supported by the Society for the Relief of Disabled Children, Hong, Kong. For the collection in Germany (Andreas Chiocchetti and Christine M. Freitag), we thank S. Lindlar, J. Heine, and H. Jelen for technical assistance, and H. Zerlaut and C. Lemler for database management. The collection was supported by Saarland University (T6 03 10 00-45 to Christine M Freitag); German Research Association DFG (Po 255/17-4 to Fritz Poustka); EC FP6-LIFESCIHEALTH (512158; AUTISM MOLGEN to Annemarie Poustka, and Fritz Poustka); BMBF ERA-NET NEURON project: EUHFAUTISM (EUHFAUTISM-01EW1105 to Christine M Freitag); Landes-Offensive zur Entwicklung wissenschaftlich-ökonomischer Exzellenz (LOEWE): Neuronal Coordination Research Focus Frankfurt (NeFF, to Christine M Freitag), and EC IMI initiative AIMS-2-TRIALS (777394-2 to Christine M Freitag). During the last 3 years, Christine M Freitag has been consultant to Desitin and Roche, receives royalties for books on ASD, ADHD, and MDD, and has been granted research funding by the European Commission (EC), Deutsche Forschungsgemeinschaft (DFG), and the German Ministry of Science and Education (BMBF). The collection in Utah was supported by R01 MH094400 (to Hilary Coon). The collection in Siena (Alessandra Renieri) was supported by the “Cell lines and DNA bank of Rett Syndrome, X-linked mental retardation and other genetic diseases”, member of the Telethon Network of Genetic Biobanks (project no. GTB12001), funded by Telethon Italy, and of the EuroBioBank network. The PAGES collection in Sweden was supported by R01 MH097849 (to J.D.B.), Supplement to R01 MH097849 (MH097849-02S1 to J.D.B.), and R01 MH097849 (to J.D.B.). The collection at University of Pittsburgh (Nancy Minshew) was supported by the Trees Charitable Trust. The collection in Brazil (Maria Rita Passos-Bueno) was supported by Fundação de apoio a pesquisa do estado de São Paulo (FAPESP)/CEPID 2013/08028-1, Conselho Nacional do Desenvolvimento Tecnológico (CNPq)466651/2014-7. For the collection in Finland (Kaija Puura), we thank The Academy of Finland (grant 286284 to T.L.); Competitive State Research Financing of the Expert Responsibility area of Tampere University Hospital (to T.L., K.P.); Signe and Ane Gyllenberg Foundation (to T.L.); Tampere University Hospital Supporting Foundation (to T.L.), European Union (The GEBACO Project no. 028696, to K.P. and M.K.), the Medical Research Fund of Tampere University Hospital (to K.P.), The Child Psychiatric Research Foundation (Finland, to M.K.) and The Emil Aaltonen Foundation (to M.K.). For the collection at UCSF (Lauren A. Weiss), we acknowledge funding sources NIH Exploratory/Developmental Research Grant Award (R21) HD065273 (to L.A.W.), Simons Foundation Autism Research Initiative (SFARI) 136720 (to L.A.W.) as well as IMHRO and UCSF-Research Evaluation and Allocation Committee (REAC) support (to L.A.W.). The collection at UIC (Edwin H. Cook) was supported by NICHD P50 HD055751 (to E.H.C), and the sequencing was funded through X01 HG007235. The Genotype-Tissue Expression (GTEx) Project was supported by the Common Fund of the Office of the Director of the National Institutes of Health, and by NCI, NHGRI, NHLBI, NIDA, NIMH, and NINDS. DDG2P data used for the analyses described in this manuscript were obtained from http://www.ebi.ac.uk/gene2phenotype/. The funders played no role in the design of the study, in the collection, analysis, and interpretation of data, or in writing the manuscript. We thank Tom Nowakowski (UCSF) for facilitating access to the single-cell gene expression data.

