## Supplementary Figures and Methods for "Large-scale exome sequencing study implicates both developmental and functional changes in the neurobiology of autism"

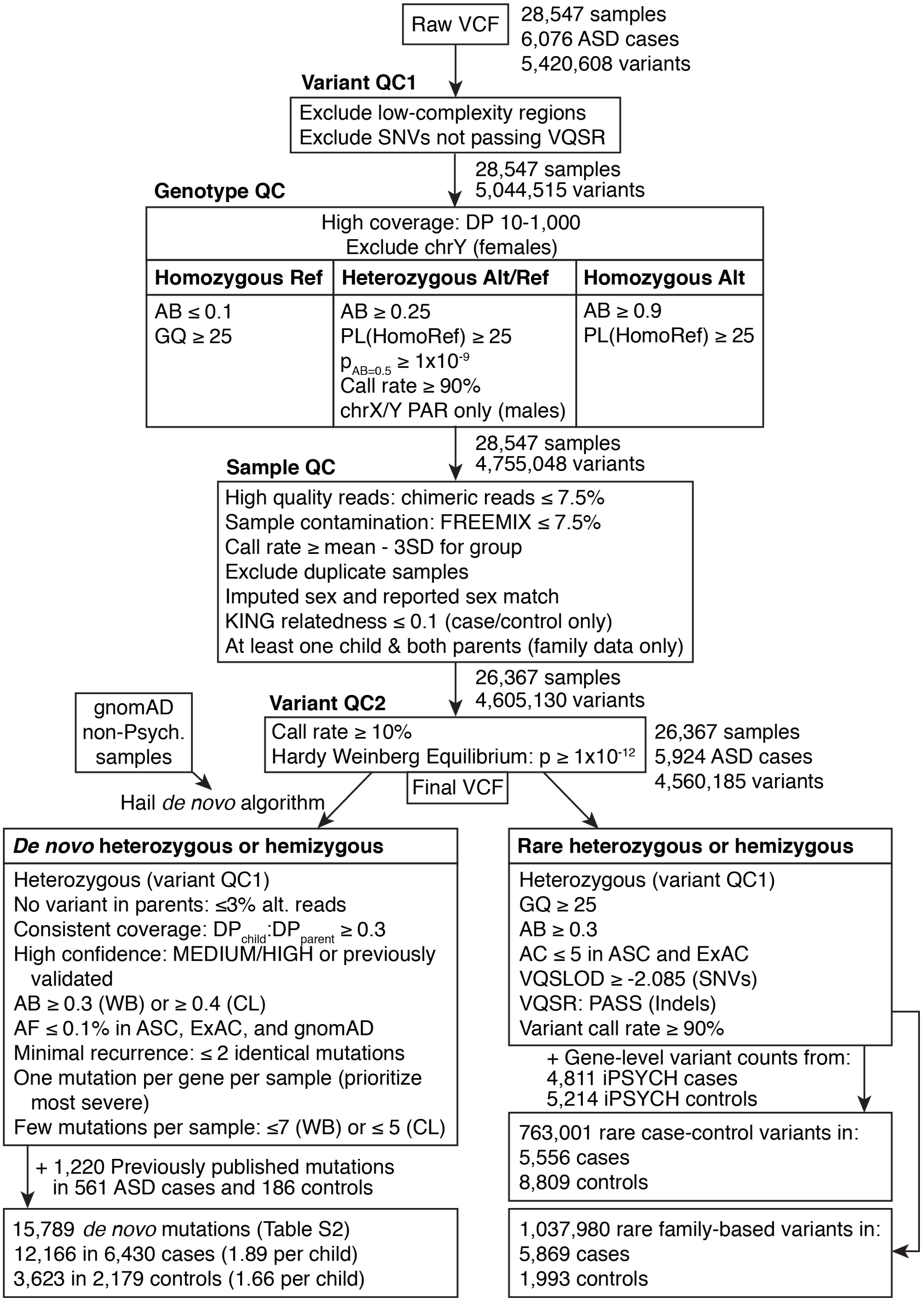


**Figure S1. Data processing for samples with read-level exome data.**

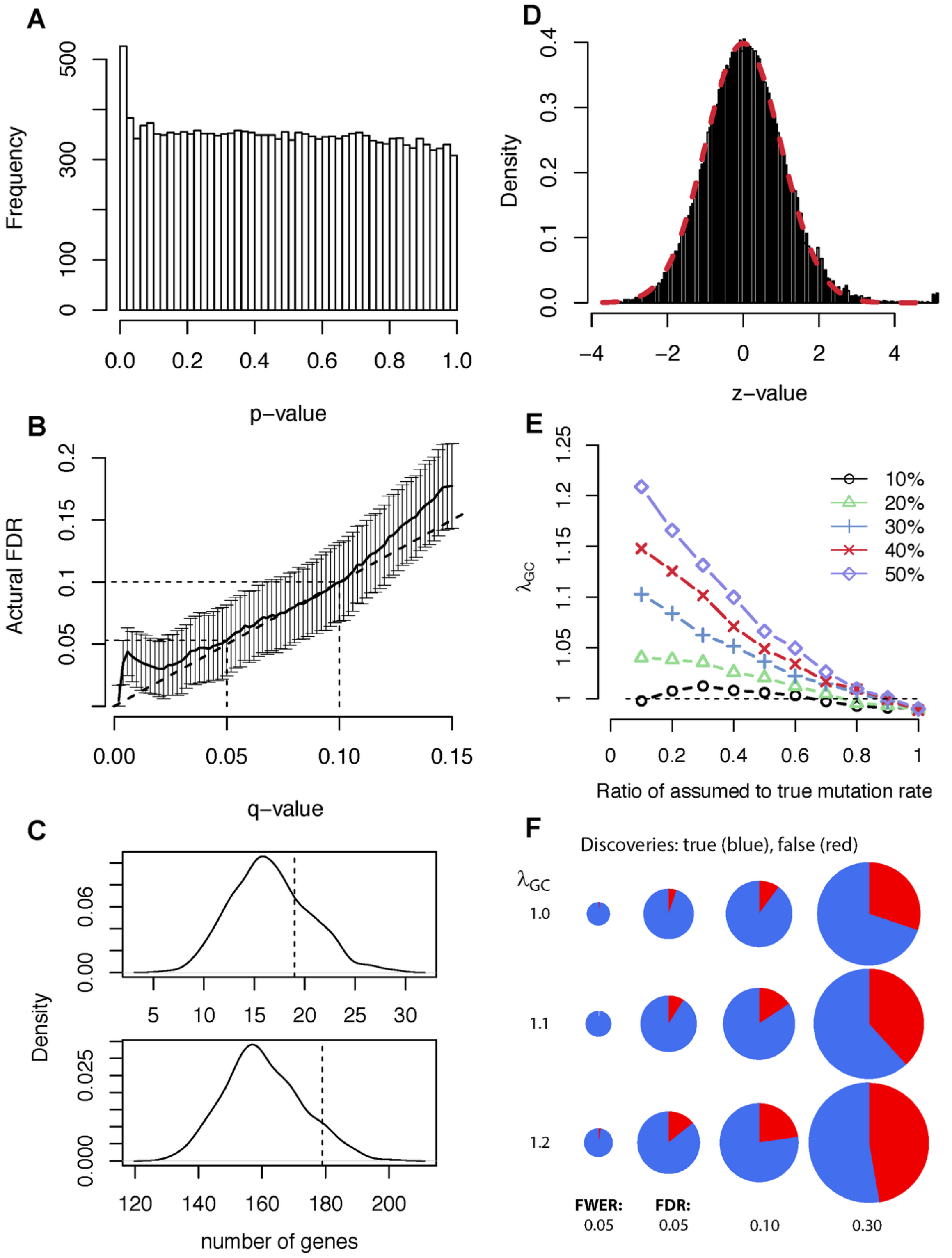


**Figure S2. Evaluating the accuracy of the TADA model. A)** Distribution of p-values from the TADA analysis of protein truncating variants (PTVs) and Mis3 (PolyPhen-2 probably damaging) variants in the data. **B)** Empirical-known signal experiments (EKSE) to evaluate the control of FDR. The bold line shows the FDR, averaged over iterations versus q-values. For these iterations, a true signal is superimposed on a common background of benign mutations. **C)** Observed vs. simulated distribution of synonymous and Mis1 (PolyPhen-2 benign) variants. The solid line shows the distribution of 500 simulation results, compared to the observed shown in the dashed line. (top) The number of genes with synonymous events ≥3. The average over simulations is 16.6, while the observed is 19. (bottom) The number of genes with Mis1 (PolyPhen-2 benign) events ≥2. The average over simulations is 160.6, while the observed is 179. **D)** To visualize potential deviations from the model in the TADA analysis of the true ASD sample, TADA p-values are converted to z-values, which are assumed to follow a standard normal density (for null genes). Here a normal density with mean zero and variance = 1.02 (in red) is superimposed on the histogram of results from the real data, illustrating how closely the data follow the assumed model (1.02 = variance = $\lambda_{GC}$ for these results). **E)** Underestimating the per-gene mutation rate could lead to false discoveries. To assess the impact of model misspecification, the genomic control factor, $\lambda_{GC}$, is plotted for various misspecifications of $\mu$. Each line corresponds to a different fraction of genes with underestimated $\mu$, and the x-axis is the magnitude of the underestimation: $\mu$-used/$\mu$-true. **F)** True discoveries (TD) and false discoveries (FD) by critical value, and for various levels of model discrepancy, with $\lambda_{GC}$ as the genomic control factor. FWER is set at the correction rate. FDR is set at 0.05, 0.10 and 0.30. The circumference of each pie chart equals the cube root of the total number of discoveries

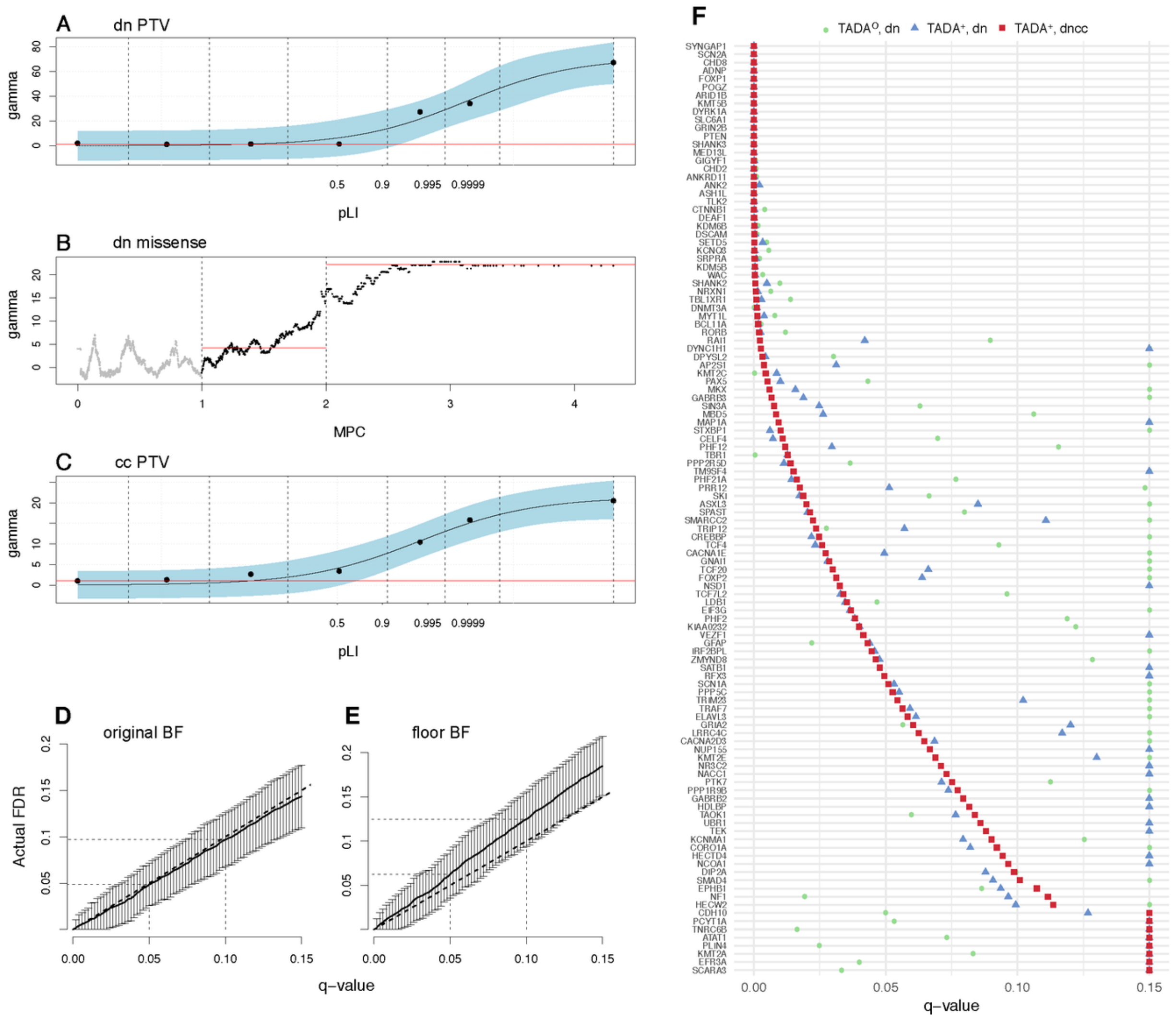


**Figure S3. Incorporation of pLI and MPC metrics into TADA. A)** The average relative risk ($\gamma$) for variants in the data. We model the relative risk for *de novo* protein truncating variants (PTVs) as a continuous function of pLI. **B)** For *de novo* missense variants, we estimate the curve using a moving window over probands' missense variants ordered by MPC score. The MPC score can be cut into three categories, and we use the last two in the TADA analysis, since $\gamma$ in the first category is close to the null value of one. **C)** For case-control PTVs we model the relative risk for *de novo* PTVs as a continuous function of pLI. **D)** The empirical FDR versus q-values, averaged over 100 simulations using the original Bayes Factors. **E)** The empirical FDR versus q-values, averaged over 100 simulations using the Bayes Factors with a lower limit. **F)** The q-values for three types of TADA analyses. We present the 114 genes with q-value less than 0.1 in at least one analysis.

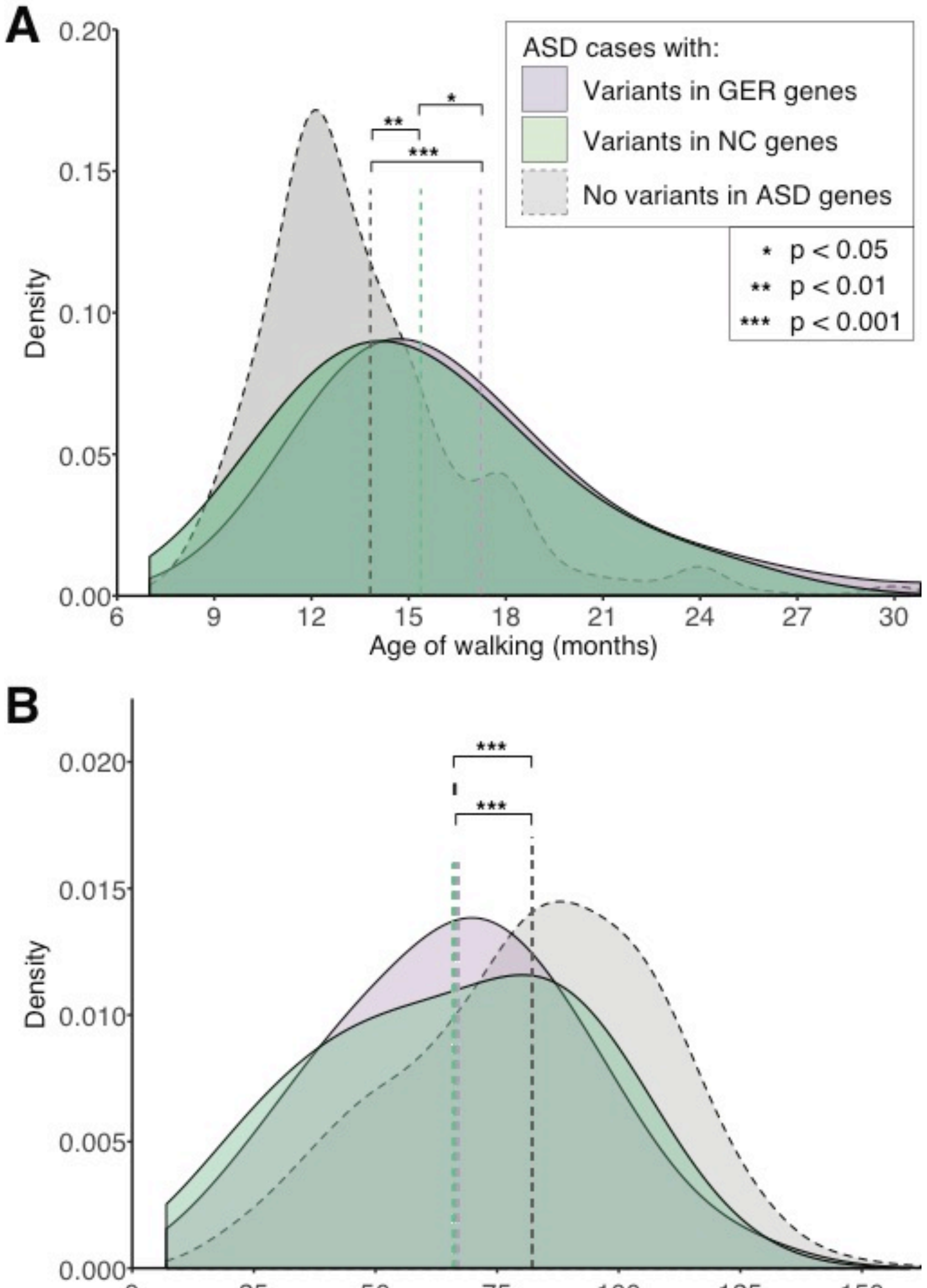


**Figure S4. Phenotypic distribution of age of walking and full-scale IQ in ASD individuals split by those carrying de novo variants in gene expression regulation (GER) genes and neuronal communication (NC) genes.** **A**) Age of walking distribution for 4456 ASD probands for whom age of walking data was available split into three groups: those carrying a *de novo* missense (MPC ≥ 1) variant or PTV in a GER gene (N=140, purple), those carrying a *de novo* missense (MPC ≥ 1) variant or PTV in an NC gene (N=71, green), and the rest of the ASD population without such a *de novo* variant in the 102 FDR<0.1 genes (N=4204, grey). **B**) Full-scale IQ (FSIQ) distribution for 4821 ASD probands for whom FSIQ was available split into three groups: those carrying a *de novo* missense (MPC $\geq$ 1) variant or PTV in a GER gene (N=159, purple), those carrying a *de novo* missense (MPC $\geq$ 1) variant or PTV in an NC gene (N=77, green), and the rest of the ASD population without such a *de novo* variant in the 102 FDR<0.1 genes (N=4542, grey).

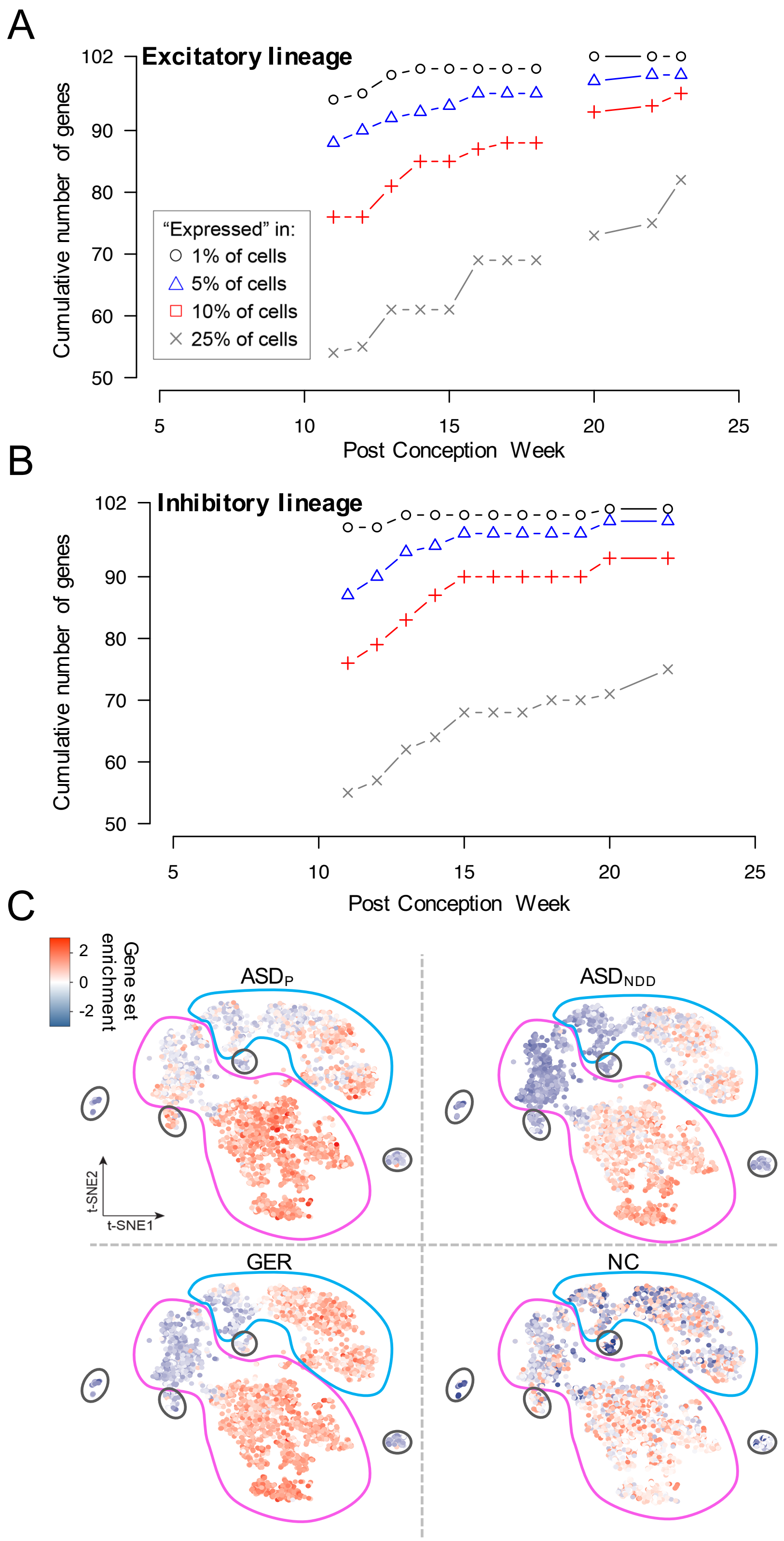


**Figure S5. Single-cell gene expression. A)** The cumulative number of ASD-associated genes expressed in RNA-seq data for different definitions of expression (0.01-0.25 of cells contain at least one transcript of the gene) and for 4,261 cells collected from human forebrain across prenatal development (Fig. 5D) in all cells from the excitatory lineage. **B)** The plot in ‘A’ is repeated for all cells in the inhibitory lineage. **C)** The enrichment of ASD associated genes in single cells arranged using tSNE (Fig. 5F) divided into the $ASD_P$, $ASD_{NDD}$, GER, and NC gene sets.

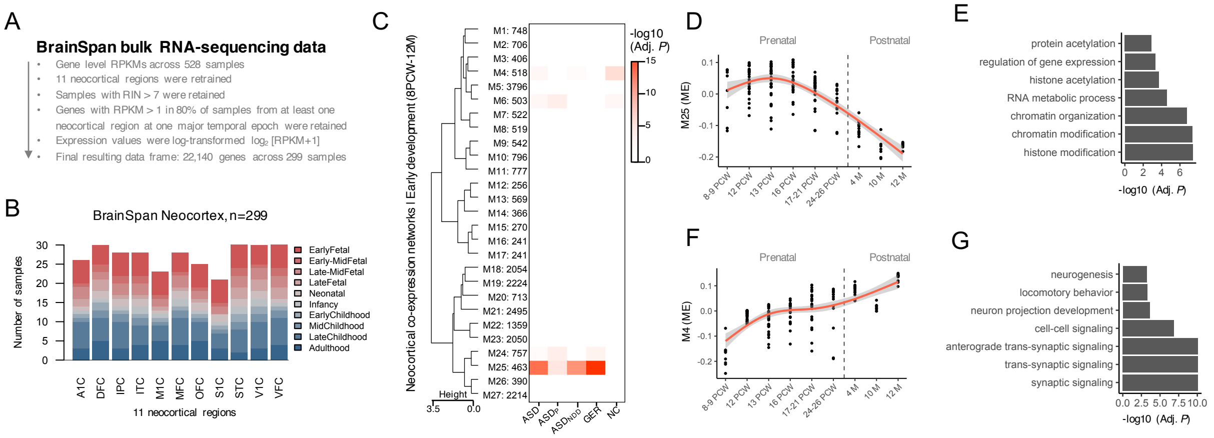


**Figure S6. Enrichment of 102 ASD genes in WGCNA modules from human cortex. A)** BrainSpan bulk RNA-sequencing data underwent extensive data quality control and data pre-processing. **B)** A total of 299 post-mortem brain samples were used to assess prenatal versus postnatal bias in gene expression patterns for each individual gene. **C)** Next, a subset of 177 high-quality samples aged 8 post-conception weeks (pcw) to 1 year of age were used to construct early developmental co-expression networks (i.e. modules). A total of 27 modules were identified, clustered by module eigengene (ME) values and assessed for enrichment for the 102 ASD risk genes. Node size indicates odds ratio and intensity signal indicates significance level. Module number and the total number of genes within each module are labeled to the right of the enrichment plot. Splines were applied to capture non-linear ME effects while ensuring patterns of gene expression are continuous across early development. **D)** Peaking in expression during mid-fetal development, module M25 was enriched for ASD and GER genes and **E)** was enriched for chromatin modification/organization and histone modification/acetylation terms. **F)** Peaking in expression during neonatal/infancy, module M4 was enriched for NC genes and **G)** was implicated in trans-synaptic signaling and cell-cell signaling.

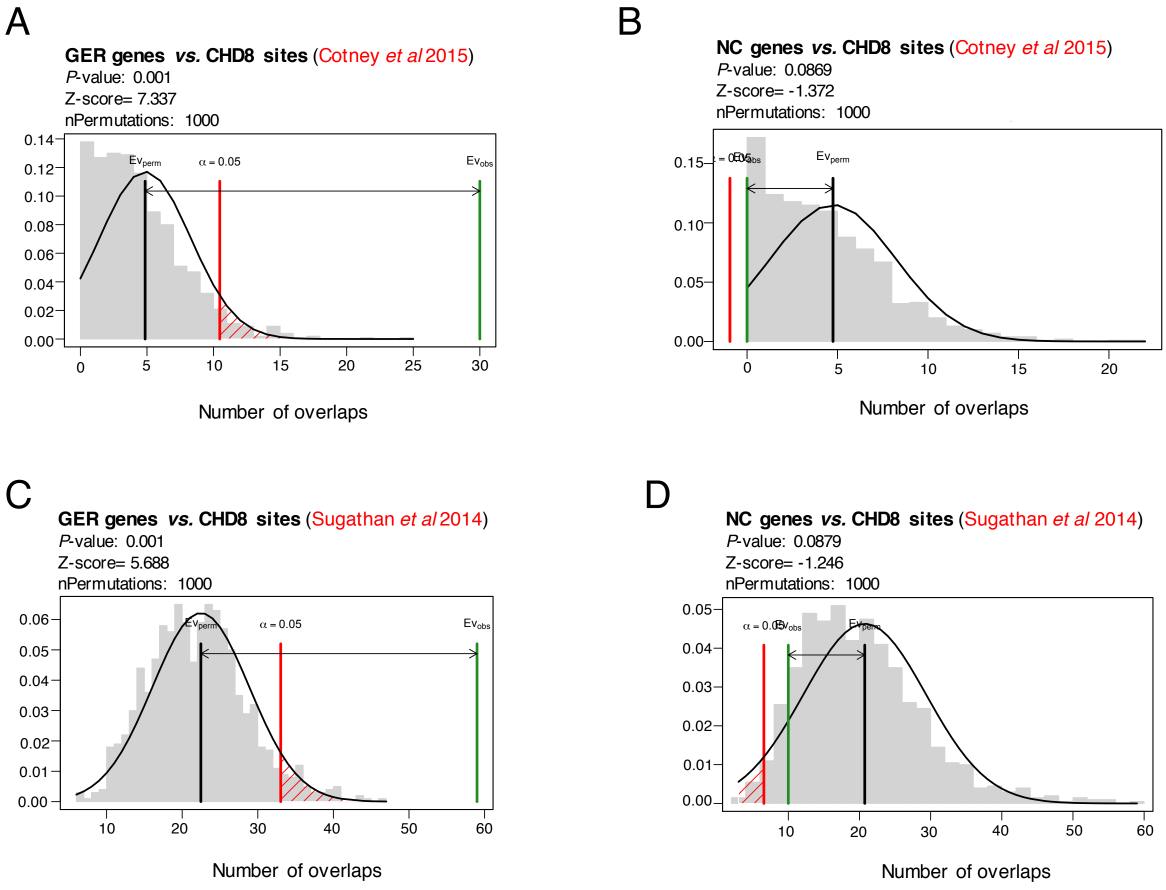


**Figure S7. CHD8 enrichment analysis.** Genomic coordinates for GER and NC genes were assessed for enrichment for human brain-specific CHD8 binding sites derived from (**A-B**) human mid-fetal brain tissue and (**C-D**) human neural progenitor cells (NPCs). The regioneR R package was used to test overlaps of genomic regions based on permutation sampling. We sampled random regions from the genome 1000 times, matching size and chromosomal distribution of the region set under study. By recomputing the overlap with CHD8 binding sites in each permutation, statistical significance of the observed overlap was computed. We observed significant enrichment for GER genes and CHD8 binding sites derived from (**A**) human midfetal brain tissue and (**C**) human NPCs. However, no significant enrichment was observed for NC genes from either study.

#### Supplementary Table List

Note: each table is its own separate file.

**Table S1. Cohorts and samples.** This file lists the cohorts that contributed samples to this study. It also contains a table of the 35,584 samples used in the analysis, as well as their cohorts of origin and other basic sample information.

**Table S2. *De novo* variants.** This table contains 15,789 predicted *de novo* variants in the 6430 individuals with ASD and 2079 individuals without ASD drawn upon in this study.

**Table S3. Liability analysis.** This file contains variant counts and calculations used in the liability analysis in Figure 1 in the main manuscript.

**Table S4. TADA FDR values.** This table lists the association data for 17,484 autosomal genes along with TADA false discovery rates. At the top are the 102 genes identified from TADA at an FDR < 0.1. Used for Figures 2b, 3a, 4a, and 4e.

**Table S5. Overlap of 102 TADA with previous literature.** This file contains a compendium of previous reports on genes involved in developmental disabilities that overlap with the 102 implicated in risk by our current study.

**Table S6. Missense variants.** This file lists *de novo* missense variants identified in DEAF1, KCNQ3, SCN1A, and SLC6A1 in this and other studies. Used to make Figures 3b, 3c, 3d, and 3e.

**Table S7. Genomic disorder (GD) loci.** This file lists GD loci and their overlap with ASD risk genes identified by TADA. Used to make Figures 3f and 3g.

**Table S8. GWAS enrichment analysis.** This table shows the gene set enrichment results from the analysis to determine if the 102 ASD-associated genes were enriched for common variants associated with ASD and genetically correlated traits. Used to make Figures 3h and 3i.

**Table S9. *De novo* variants from ascertained neurodevelopmental delay.** This table contains 8,087 published *de novo* variants in 5264 individuals ascertained for severe intellectual disability / developmental delay. Used in Figure 4a.

**Table S10. Phenotype and mutation data**. This table contains the IQ and age of walking phenotype data (when available) for the 6430 individuals with ASD and 2079 unaffected siblings used to construct Figures 4b, 4c, and 4d.

**Table S11. Functional categorization of ASD-associated genes.** This file shows specific GO terms attached to the ASD-associated genes, gives our functional classification of each gene, and outlines the evidence for the classification. Used in Figures 4e, 5a, 5c, and 6.

**Table S12. Single-cell expression analyses of the developing fetal neocortex.** This file shows the genes expressed in each cell type cluster and the enrichment of ASD genes in each cluster. Used in Figures 5d-i.

**Table S13. DAWN analysis.** This table contains full results for the DAWN (Detecting Association With Networks) analysis, which uses coexpression with known ASD-related genes to predict further ASD-related genes. Used in Figure 6a.

**Table S14. Weighted gene coexpression network analysis (WGCNA).** This file contains information related to the WGCNA analysis shown in Figure S6, including which genes are in which coexpression module and the results of gene set enrichment analyses within each module.

**Table S15. Protein-protein interaction (PPI) analysis.** This file lists the protein-protein interactions depicted in in Figure 6b. It also contains results for significance tests of module density and interaction with known DD/ID genes for the different gene sets used in this study.

**Table S16. Transcription regulator analysis.** This table lists the interactions displayed in Figure 6c.

### STAR Methods

#### CONTACT FOR REAGENT AND RESOURCE SHARING

#### EXPERIMENTAL MODEL AND SUBJECT DETAILS

The ASC was formed on the basis of prospective (i.e., prepublication) data sharing in 2010 and rapidly expanded to include about 50 groups involved in large-scale whole exome (and now whole genome) sequencing in ASD. At the outset, the ASC aggregated data from ASC sites, but over the past few years it has also been able to sequence samples at the Broad Institute / Massachusetts General Hospital through the Broad Center for Common Disease Genomics (UM1HG008895, Mark Daly, PI).

The ASC and ASC sites are approved by appropriate Institutional Review Boards or Ethical Committees. The individual studies that contribute to the ASC may have directly ascertained and interviewed human clinical subjects, along with controls, in accord with the ethical principles and practices of modern biomedical research. In contrast to contributing sites, the ASC as a group has a different relationship to human subjects. The ASC has no direct contact with the research subjects and no identifying information is provided by the primary sites to the ASC. From the perspective of the ASC, all data is de-identified and effectively anonymized.

For samples provided to the ASC sequencing site (Broad-MGH through their funding from NHGRI), a copy of the consent(s) are reviewed before samples are accepted, and a signed IRB Data Use Letter (DUL) must be received before samples are accepted. The DUL not only confirms that the samples can be shared, but it also provides the critical document for upload into dbGaP and NDAR. In this way, there are no delays in data sharing.

The iPSYCH study was approved by the Regional Scientific Ethics Committee in Denmark and the Danish Data Protection Agency.

#### QUANTIFICATION AND STATISTICAL ANALYSIS

##### Samples

The Autism Sequencing Consortium is a large-scale genomic consortium collecting and sequencing cohorts worldwide (Buxbaum et al., 2012). The analysis presented here drew from 35,584 samples collected from 32 distinct sample sets. These include cohorts sequenced by the Autism Sequencing Consortium (ASC) and published in our first (De Rubeis et al., 2014) or second study (Lim et al., 2017) (Germany, Japan, PAGES, Pittsburgh, Seaver, Spain, TASC, and UCSF), as well as new collections (Boston, Brazil, CHARGE, Chicago, Hong Kong, Miami, Portugal, Rome, Siena, Turin, UC Irvine, and Utah), with a total of 6,197 newly collected and sequenced samples included in our final analysis. We also sequenced samples from the Autism Genetic Resource Exchange (AGRE), the Boston Autism Consortium, two sites in Finland, and Swedish controls from epidemiological studies in schizophrenia and bipolar disorder. We imported exome sequence data from the Simons Simplex Collection (Iossifov et al., 2014), as well as an unpublished Norwegian cohort, and included them in our dataset alongside ASC-sequenced samples. In addition, we incorporated published *de novo* variants from the UK10K consortium, the University of Pennsylvania, Vanderbilt University, and a collection of samples from the Middle East. Finally, we integrated gene-level variant counts from autism cases and matched controls from the iPSYCH research initiative (Satterstrom et al., 2018). A description of each cohort, with the number of samples sequenced and the number used in our analyses, its ascertainment and diagnostic strategy, and associated references is found in Table S1.

The bulk of new ASC samples were sequenced at the Broad Institute on Illumina HiSeq sequencers using the Illumina Nextera exome capture kit. The remainder were sequenced at three other sites: the University of California, San Franciso (N=495), the Sanger Institute (N=443), and Johns Hopkins University (N=302), all using similar methods. Each sample's sequencing reads were aggregated into a BAM file and processed through a pipeline based on the Picard set of software tools. The BWA aligner mapped reads onto the human genome build 37 (hg19). Single nucleotide polymorphism (SNPs) and insertions / deletions (indels) were jointly called across all samples using Genome Analysis Toolkit (GATK; Van der Auwera et al., 2013) HaplotypeCaller package version 3.4. Variant call accuracy was estimated using the GATK Variant Quality Score Recalibration (VQSR) approach. The VCF file was produced by the Broad sequencing and calling pipeline with GATK version 3.4 (g3c929b0) and was itself VCF format v4.1.

**Dataset QC**

The VCF file, containing approximately 29,000 exomes, was loaded into Hail 0.1 (http://hail.is; https://github.com/hail-is/hail) to perform basic quality control steps. Multi-allelic sites were split into bi-allelic sites and each variant was then annotated with the Variant Effect Predictor (VEP, McLaren et al., 2016) by prioritizing coding canonical transcripts. VEP assigned properties such as gene name and consequence to each variant.

To check the accuracy of reported pedigree information, relatedness was calculated between each pair of samples using Hail's ibd() function and sex was imputed for each sample using Hail's impute_sex() function. The relatedness values were input into the program PRIMUS (Staples et al., 2014), which inferred pedigree structure for every related group of samples. Combined with the imputed sex, these inferred pedigrees were compared to reported pedigrees and checked for discrepancies. Obvious errors in reporting were fixed (e.g., swapped mother and father labels in the same family, or swapped parent/proband labels in the same trio), and samples with a discrepancy that could not be resolved (~200) were dropped. Parents without a child in the dataset (~250) were also dropped, resulting in 28,547 samples and 5,420,608 variants.

During a first round of variant quality control (QC), low-complexity regions were removed (110,963 variants), as were SNPs that failed variant quality score recalibration (VQSR, 265,130 variants), leaving 5,044,515 variants. For genotype QC, several genotype filters were applied: we filtered calls with a depth less than 10 or greater than 1000; for homozygous reference calls, we filtered genotypes with less than 90% of the read depth supporting the reference allele or with a genotype quality less than 25; for homozygous variant calls, we filtered genotypes with less than 90% of the read depth supporting the alternate allele or with a phred-scaled likelihood (PL) of being homozygous reference less than 25; and for heterozygous calls, we filtered genotypes with less than 90% of the read depth supporting either the reference or alternate allele, with a PL of being homozygous reference less than 25, with less than 25% of the read depth supporting the alternate allele (i.e. an allele balance less than 0.25), or with a probability of the allele balance (calculated from a binomial distribution centered on 0.5) less than $1x10^{-9}$. We additionally filtered any heterozygous call in the X or Y non-pseudoautosomal regions in a sample that imputed as male. For samples imputed as female, calls from the Y chromosome were removed. After applying these filters and removing sites that were no longer variant, the dataset contained 28,547 samples and 4,755,048 variants.

Next, we applied sample quality control filters, removing samples with estimated contamination levels >7.5% (20 samples) or chimeric reads >7.5% (121 samples). Stratifying samples into 18 different groups (by exome capture/year/cohort/sequencing center), samples were filtered if their call rate was greater than 3 standard deviations from the group mean (300 samples). Duplicate samples were then removed (761 samples), as were samples for which the imputed sex did not match the reported sex (59 samples). Following these sample filters, family structures were re-evaluated: if one or more parents of a proband had been filtered, the proband was reclassified as a case and the remaining parent (if any) was dropped; if the proband had an unaffected sibling, the sibling was kept as a "sibling of case;" if one or more parents were filtered and no proband remained, then data for remaining family members were dropped; second degree or greater relatives of probands were also dropped. After applying these rules, the dataset contained 5833 complete families, with 5924 affected probands, 2007 unaffected offspring, 5834 fathers, and 5833 mothers (one family contained two probands, two fathers, and one mother).

The dataset also contained 2388 cases, 106 siblings of cases, and 4324 controls, none of whom were part of a complete trio. From these categories, we filtered one of each related pair of samples (although each case was allowed to keep 1 sibling in the event this became interesting for future study). We defined related samples as a pair of samples with a KING (Manichaikul et al., 2010) kinship value of 0.1 or greater, approximately corresponding to a PI_HAT value of 0.2 or greater. Following this filtering, the dataset contained 2353 cases, 100 siblings of cases, and 4316 controls, for a total of 26,367 samples.

After filtering sites that were no longer variant, there were 4,605,130 variants. For a second round of variant QC, variants with call rate <10% (17,083 variants) or a Hardy-Weinberg equilibrium p-value less than $1x10^{-12}$ (27,862 variants) were filtered, leaving 26,367 samples and 4,560,185 variants. This dataset was then used as the starting point for the *de novo*, inherited, and case-control workflows. Most of the remaining samples were ultimately used in our TADA analysis, but some were subject to additional filtration during these workflows.

#### Tallying of variant classes

##### *De novo variation*

*De novo* variants were called from the 26,367-sample dataset described above, including 5924 affected probands and 2007 unaffected offspring. After filtering any genotype with a GQ < 25, *de novo* variants were called using the de_novo() function of Hail 0.1, which implements the caller used in previous ASC work (https://github.com/ksamocha/de_novo_scripts). Population allele frequencies for variants were obtained from the non-psychiatric subset of gnomAD (http://gnomad.broadinstitute.org/), and these frequencies were used as the input priors. As additional parameters, parents' homozygous reference genotypes were required to have no more than 3% of reads supporting the alternate allele, children's heterozygous calls were required to have at least 30% of reads supporting the alternate allele, and the ratio of child read depth to parental read depth was required to be at least 0.3.

This process identified 44,562 *de novo* variants (26,577 distinct variants) in the 7931 children in the dataset. Of the 7931 children, 519 were part of a whole genome sequencing project (Werling et al., 2018), and we added 168 *de novo* variants called in these samples from the whole genome sequencing that were not called in the exome sequencing. We also incorporated 338 previously published and validated *de novo* variants in our samples that were not identified by our caller (Kosmicki et al., 2017). Thus, in total, we had 45,068 *de novo* variants (27,083 distinct variants) in 7931 children. For QC on the *de novo* variants, we retained variants if they were high confidence as indicated by the calling algorithm, medium confidence and a singleton in the dataset, or previously experimentally validated (removed 20,862 calls). To filter calls stemming from cell line artifacts, an allele balance of at least 0.4 was required for samples from immortalized cell lines (773 probands and 40 siblings) (removed 2171 calls). Next, a call was removed if it had an allele frequency >0.1% in our dataset, in ExAC (r0.3, non-psychiatric subset, http://exac.broadinstitute.org/), or in gnomAD (non-psychiatric subset) (removed 5068 calls). Calls were removed if they appeared more than twice (removed 403 calls) and were then limited to one variant per person per gene (removed 570 calls), retaining calls with the most severe consequence when selecting which one to keep. Finally, samples whose DNA source was whole blood or saliva were dropped if they had more than 7 coding variants (removed 20/5143 probands and 13/1967 unaffected children), and samples whose DNA source was immortalized cell lines were dropped if they had more than 5 coding calls (filtered 35/773 probands and 1/40 unaffected children). Retained were 14,569 *de novo* variant calls from 5869 probands and 1993 unaffected children. To maximize power, we supplemented this set with 933 and 287 published *de novo* variants in 561 probands and 186 siblings (De Rubeis et al., 2014; Sanders et al., 2015; Kosmicki et al., 2017), respectively, for whom original sequence data were not available.

*Inherited variation*

QC for inherited variants began with the dataset of 26,367 samples and 4,560,185 variants. Any genotype call with a GQ < 25 was removed and heterozygous genotypes were required to have an allele balance ≥ 0.3. Variants were required to have a call rate ≥90%, insertions and deletions were required to pass VQSR, and SNPs were required to have a VQSLOD (variant quality score log odds) ≥ -2.085. The VQSLOD threshold for SNPs was determined by identifying the threshold at which synonymous variants with an allele count of 1 amongst parents in the dataset were transmitted to the child 50% of the time, as described previously (Lek et al., 2016; Kosmicki et al., 2017). Protein-truncating variants were required to be high confidence ("HC") by the LOFTEE plugin for VEP and to have no LOFTEE flags other than "SINGLE_EXON".

For purposes of gene-level counts, variants were tallied in the 5869 probands and 1993 unaffected children who passed *de novo* QC. Variants were required to have an allele count ≤ 5 in the combined parents, cases, and controls (18,153 people) in our dataset, as well as an allele count ≤ 5 in the non-psychiatric subset of ExAC.

*Case-control variation*

Variants in ASC cases and controls were QC'd in the same way as inherited variants. For purposes of gene-level counts, variants were again required to have an allele count ≤ 5 in the 18,153 combined parents, cases, and controls in the dataset, as well as an allele count ≤ 5 in the non-psychiatric subset of ExAC.

To ensure well-matched cases and controls, probable ancestry was calculated by merging our raw dataset with genotypes from the 1000 Genomes Project and conducting principal components analysis (PCA) in Hail on a set of ~5000 common SNPs. A naive Bayes classifier was trained (using the naiveBayes function from the R package e1071) on the 1000 Genomes samples labeled as either European or East Asian and used to predict which of our samples fell into those populations. Synonymous rates were well-matched between cases and controls from the Swedish contributing site which were classified European (745 cases and 3595 controls), as well as between cases and controls from the Japanese contributing site which were classified East Asian (196 cases and 298 controls). For inclusion in TADA, we counted variants from the 4340 Swedish samples. Overall variant rates were higher in the Japanese samples than the Swedish samples, possibly because our filtering was based on allele counts in ExAC, and ExAC has less representation from East Asian samples than European ones.

**Analysis of variant classes**

To model a qualitative trait—in this case, the presence or absence of ASD—using standard quantitative genetics concepts, we imagine that there is an unobserved, normally distributed variable called "liability" that determines whether or not an individual is diagnosed with ASD. We assume that liability, $L$, has mean 0 and variance 1 in the general population. Individuals with $L$ greater than some threshold $t$ are diagnosed with ASD and individuals with $L < t$ are considered "typical". Under this model, the prevalence difference between males and females is viewed as a difference in thresholds for males and females. For a male to be diagnosed with ASD, his liability must be larger than $t_m$. For a female to be diagnosed with ASD her liability must be larger than $t_f$. Since ASD is more common in males than females, we conclude that $t_m < t_f$. For all that follows we will assume that the prevalence of ASD, $\Psi_m$, is 1 in 42 in males (implying $t_m \sim 1.98$), and the prevalence of ASD is 1 in 189 females, $\Psi_f$ (implying $t_f \sim 2.56$). We model ASD+ID similarly, but with lower prevalence than all ASD (male prevalence 0.00499, and female 0.00138).

When considering the effects of individual alleles on liability, we employ an elaboration to the standard quantitative genetics model, which is sometimes called the "mixed model of inheritance". We assume that individual alleles make additive contributions to liability, so that for some allele, $A_1$, individuals with 0 copies of the allele have mean $-\mu$, variance 1 liability, but individuals with 1 copy have mean $\alpha - \mu$, variance 1, and individuals with 2 copies have mean $2\alpha - \mu$, variance 1 liability. Assuming Hardy-Weinberg equilibrium for genotypes, and the frequency of $A_1$ equaling $p$, $\mu = 2\alpha p^2 + \alpha 2pq = 2\alpha p$. Here $\mu$ is a normalizing factor to ensure the overall population has mean liability 0.

For several of our analyses we are interested in the effect, $\alpha$, for variants of a particular type in a collection of genes, for instance *de novo* PTVs in genes with pLI scores > 0.9. If a variant is individually exceptionally rare, we have virtually no power to estimate its individual effect size, but over a large collection of such variants average properties are estimable. To do so, we model the entire collection of variants as if there were a single allele with frequency equal to the sum of the individual variant frequencies. This approach makes little sense for common variants, but for sufficiently rare variants, where single individuals seldom harbor more than one, this is a reasonable and helpful approximation. For some variant types, however, such as silent variants, the count of alleles can be substantial. For this reason, rather than standardize by $2N$, where $N$ is the number of subjects, we standardize by $2NM$, where $M = 17{,}484$ is the number of genes analyzed herein. This standardization has no material impact on calculations of parameters of interest. To distinguish between cases and controls, we write $N_{ca}$ and $N_{co}$ respectively.

Thus, for each type of variant we are interested in studying, *de novo* PTV mutations, say, we count the number of observations of this class of variant in cases (our probands in trios), and the number of observations of this class of variant in controls (our siblings in trios). For a given type of variant, $V$, we call $Pr\{V|D\}$ the frequency of this type of variant in cases (observed number of variants divided by $2NM$), and $Pr\{V|\neg D\}$ the corresponding value in controls. We make these calculations separately in males and females, which we denote as $Pr\{V_m|D_m\}$, $Pr\{V_f|D_f\}$, $Pr\{V_m|\neg D_m\}$, and $Pr\{V_f|\neg D_f\}$ where the $m$ and $f$ subscripts distinguish male and females. The overall frequency of the variant class can be found by

$$Pr\{V_g\} = Pr\{V_g|D_g\}\Psi_g + Pr\{V_g|\neg D_g\}(1 - \Psi_g)$$

where $g$ can be either $f$ or $m$, for females and males, respectively. From this the Penetrance (probability of disease given variant) of the variant class can be found immediately by Bayes rule

$$Pr\{D_g|V_g\} = \frac{Pr\{V_g|D_g\}\Psi_g}{Pr\{V_g\}}.$$

To find the average effect, $\alpha_{V_g}$, of this variant class we note

$$Pr\{D_g|V_g\} = \int_{t_g - \alpha_{V_g}}^{\infty} \frac{1}{\sqrt{2\pi}} e^{-\frac{x^2}{2}} dx.$$

Thus, we can find the effect size by inverting a standard normal cumulative distribution, $\Phi(x)$

$$\alpha_{V_g} = t_g - \Phi^{-1}(1 - Pr\{D_g|V_g\}).$$

Empirically, the relative risk for the variant type is calculated as $Pr\{V_m|D\}/Pr\{V|\neg D\}$ for the contrast of cases versus controls. To assess whether or not there is any difference in this variant class between cases and controls, we perform an exact Binomial Test on the underlying observed counts, where the probability of success is given by $N_{ca}/(N_{ca} + N_{co})$. The odds ratio is computed from four observations, the number of variants of the risk class in cases, $a$; the number of variants of the risk class in controls, $b$; the number of alleles not in the risk class in cases $2N_{ca}M - a$; and the parallel calculation for controls, $2N_{co}M - b$.

To estimate a confidence interval of $\alpha_{V_g}$, we note that in a very formal sense $\alpha_{V_g}$ is the average effect on the liability scale of the variant. Were we able to observe those effects directly, we could have calculated the observed mean and standard error of those effects. Because we cannot observe liability directly here, we infer the standard error of $\alpha_{V_g}$ by the following procedure: map the p-value from the Binomial test, described above, onto an equivalent z-value from the normal distribution, $z$; then $\alpha_{V_g}/z$ is a reasonable estimator for the standard error of the estimator for $\alpha_{V_g}$.

For Table S3, calculations for "All Genes" and for "Other Genes" were performed separately for males and females and also separately for the PAGES and DBS samples. Inherited analysis calculations were also separated by male / female and by proband / sibling. To combine effects between males and females, we took inverse-variance weighted averages of male and female effect sizes. We performed analogous calculations for the populations of case-control samples. For these calculations for the 102 ASD genes, however, because the counts of events were often small, we combined data over males and females and over PAGES and DBS samples to compute overall parameters (i.e., performed mega- versus meta-analysis). When parameters could not be estimated, this is noted as NA.

#### Transmission And De novo Association test (TADA)

*Background*

Published analyses of WES data using TADA have evaluated two categories of rare variation, namely protein truncating variants (PTV; i.e., frameshift, stop gained, splice site acceptor or donor mutation) and probably damaging missense according to PolyPhen-2 (Mis3) (Adzhubei et al., 2010), in the context of three categories of inheritance pattern: *de novo*, inherited, and case-control. TADA requires a mutational model (Sanders et al., 2012; Neale et al., 2012; Samocha et al., 2014) which accounts for gene size and sequence composition to obtain an expectation for mutations per gene, given sample size. It treats all PTV mutations within a gene as equivalent, although their impact on risk is allowed to vary across genes and inheritance patterns (likewise for Mis3). TADA first computes a gene-specific Bayes factor for each mutation category and inheritance pattern, and then it multiplies these Bayes factors to generate a statistic that summarizes all evidence of association for each gene. The total Bayes factor is finally converted to a q-value to control FDR (De Rubeis et al., 2014). As a Bayesian model, TADA requires prior parameters or hyperparameters, namely the fraction of genes in the genome affecting risk, thus far taken to be 0.05, and $\gamma$, the relative risk for a particular mutation category. See He et al. (2013) for estimators.

*Evaluating TADA and False Discovery Rate (FDR)*

For downstream analysis it is critical to ensure reliable performance of TADA so that risk gene lists, such as those with FDR $< 10\%$, are properly calibrated. Such guarantees are straightforward to prove in many settings (Efron, 2012). In the WES setting, however, and especially for the relatively discrete counts of *de novo* events, a demonstration that the FDR rate holds is warranted. It is worth noting that, even though there are many genes that contain no mutations, the mutation rate is gene-specific and varies with gene length. Consequently, with the exception of the genes with a signal, the p-values from the TADA analysis of PTV and Mis3 mutations are almost uniformly distributed (Figure S2A).

To evaluate the validity of the FDR framework in the context of TADA analysis, we conduct "empirical-known signal experiments" (EKSE). The idea is to perform TADA analyses in which the true signal is known *a priori*. To make the simulation as real as possible, it is performed using real *de novo* mutation counts as a base. These mutations are chosen to carry no detectable signal (i.e., mimicking the null distribution because they are believed to be non-functional). Simulated signals for association are then generated for randomly selected genes. Once the data are generated, TADA is used to analyze them and the resulting FDR and other features of the method are examined.

*EKSE Simulations to Assess the Properties of FDR.* For these empirically-known signal experiments, we let synonymous variants play the role of Mis3 (denoted as Mis3new) and Mis1 play the role of PTV (denoted as PTVnew). Signals are layered onto genes that are randomly chosen. Below is the detailed procedure:

1. Divide all 17,484 genes into 20 bins of equal size. Let $b = 1, \dots, 20$.
2. For each of the 20 bins, iteratively generate a signal for all genes in the bin; the remaining 19 bins, with no signal, represent the null genes. The extra signal in the $i$th gene for both new kinds of *de novo* variants is simulated using $X_i|\gamma_i \sim \text{Poisson}(2\mu_i(\gamma_i - 1)N)$, where $\gamma_i \sim \text{Gamma}(\gamma, \beta)$. The hyperparameters are selected to yield signals similar to the real data: $\beta = 0.2$ and $\gamma$ is set to be 2.4 and 5.4 for Mis3new and PTVnew respectively, and $N = 6430$. The "-1" is to account for the observed *de novo* variants already included from the real data. The simulated *de novo* events are added to the observed Mis3new and PTVnew to create each of the 20 data sets.
3. Perform TADA analysis for each of the 20 datasets.
4. Display the resulting q-FDR curves for $b = 1, \dots, 20$, and q-FDR averaged over $b$.

*Pure Simulations to Assess the Properties of FDR.* This simulation is closely related to EKSE. The only difference is that the null mutations are generated randomly from a multinomial distribution instead of adopted directly from the Syn and Mis1 variants. The procedure is described below.

1. Randomly sample a fraction of all 17,484 genes as signal genes, denoted as set $S$. We set the fraction as $\pi = 0.05$. The number of trios is $N = 6430$.

2. For both two new types of variants, Mis3new and PTVnew, the mutations of all the genes are randomly generated from a multinomial distribution, $\mathbf{X} \sim \text{Multinom}(\text{M}, \mathbf{p})$, where the probability vector $\mathbf{p}$ is proportional to $\mathbf{p} = \{\mu_i \gamma_i\}_{i=1,\dots,17484}$, where $\gamma_i \sim \text{Gamma}(\gamma, \beta)$ if $i \in S$, otherwise equals 1. The total number of mutations is $M = 2N \sum_{i=1}^{17484} \mu_i \gamma_i$. The mutation rates of Mis3new are taken from Syn, and the mutation rates of PTVnew are taken from Mis1. The hyperparameters $\gamma, \beta$ are set to be the same as in EKSE.
3. Perform TADA analysis on the two generated types of variants. Display the resulting q-FDR curve.
4. Repeat steps 1-3 100 times.

Figure S2B shows the averaged actual FDR versus the q-value over the 20 EKSE experiments. The error bars are obtained from the pure simulation. For $q < 0.1$ the average curve follows the diagonal line (roughly), which indicates that the actual FDR is well controlled in the region of primary interest. We do detect a slight bump in the actual FDR for $q > 0.1$. To understand this deviation we compared the observed counts for synonymous (Mis3new) and Mis1 (PTVnew) to simulated counts generated from the model.

The distribution of the number of genes with synonymous counts ≥3 and Mis1 counts ≥2 is contrasted with the observed counts (Figure S2C). The contrasts show that there is a slight excess of multiple hits in the observed counts compared to the model. Adding counts of synonymous and Mis1 mutations we obtain a single distribution of mutations per gene and find that there is an excess of counts of 0, 2, 3, and >3 and a relative lack of counts of 1; overall the counts are fairly similar, but they differ significantly from expectations (chi-square p-value = 0.012). The 8 null genes with the strongest TADA signal are *GNS, LRRFIP1, GALC, GRN, MYH9, FOXK2, AP1B1*, and *UNC45B*, and these are the genes that contribute to the bump in the FDR. However, none of these genes are significant ($q < 0.1$) in the EKSE analysis or in the actual data analysis of Mis3 and PTV mutations. From this EKSE experiment we conclude that the TADA model does not perfectly capture reality and the actual FDR deviates slightly from reported value for values of $q > 0.1$. This deviation is likely due to inexact estimates of the per gene mutation rate.

TADA relies on a mutation rate model for genes, which is an estimated quantity. Hence we evaluate the impact of misspecification of mutation rates. To quantify the deviation from the expected null distribution due to mutation rate misspecification, we use the theory of genomic control (Devlin & Roeder, 1999), specifically estimating the inflation factor $\lambda_{GC}$. In this experiment we randomly select 10-50% of genes and artificially make the nominal mutation rates increasingly lower than their true mutation rates. This will make the observed mutation count larger than the expected count for a subset of genes. The result is that test statistics for association will tend to be increased for some genes, and the larger the discrepancy, the larger the set of test statistics that do not follow the expected null distribution. The genomic control factor, based on the z-statistics from the TADA analysis (Figure S2D), quantifies this inflation. As expected, the genomic control factor increases as more genes are analyzed with lower nominal than true mutation rates (Figure S2E). The inflation for $\lambda_{GC}$ is modest, however, even for these fairly notable misspecifications of the mutation rates.

Because TADA is a Bayesian method it is more natural to use FDR than a Family-Wise Error Rate (FWER) cutoff to determine significance. In this gene discovery setting it is informative to compare the numbers of true discoveries (TD), false discoveries (FD), and FDR for different p-value and FDR thresholds and to examine the impact of model mis-specifications on FDR (Figure S2F). We measure discrepancies via the genomic control factor ($\lambda_{GC}$). We simulate the Z-value of 20,000 genes, 5% with a signal from $N(\mu, \lambda_{GC})$ and 95% from the null $N(0, \lambda_{GC})$, where $\lambda_{GC}$ varies from 1 to 1.2. The value of $\mu$ is chosen to be 2 to approximately mimic the real data. Based on 1,000 replications, we calculate the average TD, FD, and FDR for a Bonferroni adjusted p-value threshold and different FDR thresholds. As expected, FWER has considerably fewer FD but also notably fewer TD than FDR, and the observed FDR is well calibrated when $\lambda_{GC} = 1$ (Figure S2F). (For $\lambda_{GC} = 1$, TD = 5, 52, 113, 334, and FD = 0.1, 3, 13, and 144 for the four thresholds examined. In each case the error rate is controlled at the expected rate.) However, as $\lambda_{GC}$ increases the actual FDR increases rapidly, especially for larger q-values. In contrast, FWER is fairly well controlled even for model discrepancies.

*A more powerful TADA model*

TADA requires input of several parameters, most notably the relative risk, $\gamma$. To estimate the relative risk for a category of mutations, we use the burden-relative risk relationship derived in He et al. (2013): $\gamma = 1 + (\lambda - 1)/\pi$, where $\pi = 0.05$ is the estimated fraction of risk genes and the burden $\lambda$ is calculated by comparing mutation counts in probands and unaffected siblings. Because differences in sequencing depths and variant calling procedures may lead to systematic differences in mutation rates, we normalize the counts using synonymous mutations counts. Let $x$ and $S$ be the number of mutations in the category of interest and compare the counts in cases (cs) and controls (cn) as $\lambda = (x_{cs}S_{cn})/(x_{cn}S_{cs})$.

Previous TADA analyses (De Rubeis et al., 2014; Sanders et al., 2015) used two annotation categories, PTV and Mis3. Here we develop a more powerful version of TADA, which uses additional annotation information. For clarity we will label the original version $TADA^0$ and the refined model $TADA^+$.

Recent studies have refined our understanding of what variation is likely to be meaningful for risk in two ways. Regarding PTVs, Kosmicki et al. (2017) showed that signals carried by PTVs involve a subset of genes that are evolutionarily constrained. For these genes, the population tends to have far fewer PTVs than would be expected based on gene size, base-pair content and evolutionary models. This constraint feature of genes is embodied in pLI (the probability of being loss-of-function [aka PTV] intolerant) (Lek et al., 2016), which is a metric ranging from zero to one, with a larger pLI representing a greater dearth of PTV variation. Kosmicki et al. (2017) found that genes with pLI $> 0.9$ tend to harbor most of the ASD association signal from PTVs. In this work, we model the relative risk ($\gamma$) of *de novo* PTVs as a continuous function of pLI. Figure S3A shows $\gamma$ as a function of pLI, where the x-axis has been converted using the inverse normal transformation, but the original values of pLI are given. We create seven bins of data and fit a logistic curve to the data. The dots are the data and the black line is the fitted curve. Then we compute error bars based on the 95% prediction interval around the fitted curve. In the upcoming implementation we truncate $\gamma$ at the null value of one.

More refined information is also available for missense variants. Samocha et al. (2017) recently introduced the MPC score, a missense deleteriousness metric composed of “Missense badness”,

PolyPhen-2 (Adzhubei et al., 2010), and Constraint. This metric also uses the concept of evolutionary constraint and seeks to quantify the degree of constraint for all missense variation in the genome. To determine how MPC might be used in TADA, we compute the average relative risk ($\gamma$, the hyperparameters for TADA) for a moving window of MPC in the ASC data (Figure S3C). Using a window over probands' missense variants ordered by MPC score, and with a width of 7.5% of the variants, we obtain the curve showing the average relative risk as a function of MPC score. Three levels of $\gamma$ naturally emerge from this relationship, with the first level (MPC $< 1$) being close to marginal relative risk and two levels showing evidence for excess burden in ASD (Figure S3C). Based on the nature of these results, we chose to group missense mutations into two categories for TADA, using established thresholds of MPC (Samocha et al., 2017): $1 \leq$ MPC $< 2$ (MisA) and MPC $\geq 2$ (MisB). Note that missense variation with MPC $< 1$ is treated as benign. The relative risk for each of the two missense categories is computed directly from the data (He et al., 2013) as $\gamma_{MisA} = 4.18$ and $\gamma_{MisB} = 22.15$.

Besides the *de novo* variants, we also consider PTVs from case-control data by aggregating the iPSYCH (Danish) data and PAGES (Swedish) data. Following the same procedure as for *de novo* PTVs, within seven bins, we estimate the relative risks for the two case-control datasets separately and combine them with a precision weight. We then fit a logistic curve using the seven points to smooth $\gamma$ as a continuous function of pLI (Figure S3D). In the TADA analysis, we treat $\gamma$ of each gene as fixed for case-control data to achieve closed-form solutions and thus facilitate the computation.

These analyses define three categories of mutation potentially meaningful for risk. The gene-specific mutation rates for PTVs and missense variants have been reported previously (Lek et al., 2016), and we further estimated the mutation rates for MisA and MisB. With mutation rates and hyperparameters estimated above, the refined TADA model can be applied to the data to identify risk genes for ASD. To allow for more variability in the prior for $\gamma$, we set $\beta = 0.2$.

To resolve an emerging issue with the model's Bayes factor (BF) values, we implement a floor adjustment that imposes a lower bound of 1 on all BF. The issue is that for some genes with larger mutation rates and zero *de novo* MisB mutations, the MisB BF is $<< 1$. Multiplying this with the other evidence renders those genes not significant. Indeed, with the mutation rates provided and the high relative risk of MisB, the model clearly expected to observe at least one *de novo* MisB variant. (This happens for other categories as well, but most notably for MisB.) We think the problem is heterogeneity of genes—some genes with a *de novo* PTV just do not have MisB mutations in the data, even though these mutations are expected. It does not make sense to have the observation of no mutations drive the model. To circumvent the problem, we made a modification of the method so that BF is replaced by max(1,BF). We tested this in simulations and the size of the modified test is satisfactory (see the discussion in the next section).

$TADA^{+}$ incorporates all of the refinements delineated here. Using $TADA^{+}$, 102 genes with q-value less than 0.1 are identified, including three genes that have excessive PTV in siblings (*EIF3G*, *KDM5B*, *RAI1*). By contrast, $TADA^{0}$ identifies only 79 genes when applied to the same data. Clearly, the new relevant functional information embodied in the pLI and MPC scores improves the power of TADA by refining the model.

*Simulations to evaluate TADA*[+]. Simulations illustrate the performance of TADA[+] when applied to *de novo* mutations only. In this setting, we simulate three types of *de novo* variants: PTV, MisA, and MisB, using the mean risks and mutation rates from real data. Below is the detailed procedure.

1. Randomly select 5% of 17,484 genes as the signal genes; denote the set of signal genes as $G_S$ and the null genes as $G_N$.
2. For each signal gene $g \in G_S$, generate risk $\gamma_g$ for each three types of variants from a Gamma distribution, $\gamma_g^a \sim \mathrm{Gamma}(\bar{\gamma}_g^a, \beta)$, $a = (PTV, MisA, MisB)$, where $\beta = 0.2$ and the hyper parameters $\bar{\gamma}_g^a$s are set to match the empirical counts. Note that $\bar{\gamma}_g^{MisA}$ and $\bar{\gamma}_g^{MisB}$ are the same across all genes, but $\bar{\gamma}_g^{PTV}$ are different.
3. For the null genes $g \in G_N$, set $\gamma_g^{PTV} = \gamma_g^{MisA} = \gamma_g^{MisB} = 1$
4. For each variant, generate the counts from a Multinomial distribution, where the total number is the expected total counts $2N \sum_g \mu_g^a \gamma_g^a$, $N = 6430$, and the probability is proportional to $\{\mu_g^a \gamma_g^a\}_{g=1}^M$. The mutation rates are taken from the real data.
5. Apply TADA[+] with Bayes Factors having a lower limit (floor) of 1, and calculate the empirical FDR.
6. Repeat steps 1-5 100 times.

Figure S3F shows that the TADA[+] model controls FDR. Applying the floor principle increases the false discovery rate by a modest amount. In practice we found that there was considerable heterogeneity across genes and this adjustment was necessary.

*TADA analyses*

We explore the performance of TADA[0] and TADA[+] with three analyses:

A. TADA[0] applied to ASC2018, *de novo* only;
B. TADA[+] applied to ASC2018, *de novo* only; and
C. TADA[+] applied to ASC2018, *de novo* and case-control data.

By moving through the three analyses, we change one variable at a time and analyze the consequences. From A to B, we compare the improvements in the model by contrasting TADA[0] and TADA[+]. From B to C, we assess the impact of adding in the case-control data.

With additional data and a more powerful TADA model, we obtain substantial new discoveries. We identify 65 genes in A, 85 genes in B, and 102 genes in C. We can visualize the q-values of the three analyses for the 114 genes with q-value less than 0.1 in at least one analysis—for most genes, the q-values decrease in sequence from analysis A to B to C (Figure S3F), with the q-value of analysis C being the smallest. Twelve genes have a q-value greater than 0.1 in C but less than 0.1 in at least one other analysis; of these 12 genes, most are downgraded in analysis C because of refinements in the new TADA model (with the genes or variants, having, for instance, low pLI score or low MPC score, particularly MPC less than 1 or missing and thus not categorized as MisA or MisB).

**Defining gene groups**

Past analyses have identified two major groups of ASD-associated genes: those involved in gene expression regulation (GER) and those involved in neuronal communication (NC) (De Rubeis et al., 2014; Sanders et al., 2015). A simple gene ontology analysis with our list of 102 ASD genes replicates this finding, identifying 16 genes in category GO:0006357 "regulation of transcription from RNA polymerase II promoter" (5.7-fold enrichment, FDR=6.2x10-6) and 9 genes in category GO:0007268: "synaptic transmission" (5.0-fold enrichment, FDR=3.8x10-3). To assign genes to the GER and NC categories, we used a combination of gene ontology, gene descriptions, and primary research.

58 genes were assigned to the GER group based on one of:

1. Clear description of a role as a chromatin modifier, transcription factor, or DNA/RNA binding protein on RefSeq (O'Leary et al., 2016)
2. Evidence of role as a chromatin modifier, transcription factor, or DNA/RNA binding protein in primary research on pubmed
3. Located in the nucleus (GO:0005634) and annotated with at least two of the following gene ontology groups or their children:
   - GO:0000122:negative regulation of transcription by RNA polymerase II
   - GO:0000785:chromatin
   - GO:0000981:RNA polymerase II transcription factor activity, sequence-specific DNA binding
   - GO:0000988:transcription factor activity, protein binding
   - GO:0003682:chromatin binding
   - GO:0003700:DNA-binding transcription factor activity
   - GO:0006325:chromatin organization
   - GO:0010468:regulation of gene expression
   - GO:0045944:positive regulation of transcription by RNA polymerase II

24 genes were assigned to the GER group based on one of:

1. Clear description of a role in the synapse or regulating membrane potential on RefSeq (O'Leary et al., 2016)
2. Evidence of a role in the synapse or regulating membrane potential in primary research on pubmed.
3. Located in the cytoplasm (GO:0005737) and annotated with at least two of the following gene ontology groups or their children:
   - GO:0007267:cell-cell signaling
   - GO:0042391:regulation of membrane potential
   - GO:0045202:synapse

Of the remaining 20 genes, 9 are annotated with GO:0007010:cytoskeleton organization or child terms of this ontology term; these genes are classified as "Cytoskeleton genes". The remaining 11 genes are described as "Other". See Table S10 and Figure 4.

**<u>Comorbid phenotypes</u>**

Full-scale IQ scores were measured using several tests including, but not limited to, the Differential Ability Scales, Second Edition (Elliott, 2007); the Mullen Scales of Early Learning (Mullen, 1995); the Wechsler Intelligence Scale for Children (Wechsler, 1992); and the Wechsler Abbreviated Scale of Intelligence (Wechsler, 1999). The full-scale IQ estimates were taken from the full-scale deviation IQ variable when available and full-scale ratio IQ when it was not (Chaste et al., 2015). Full-scale IQ is normally distributed with a mean of 100 and a standard deviation of 15. We defined intellectual disability to be if a subject met one of the following conditions: a full-scale IQ (FSIQ) < 70 (i.e., two standard deviations below the mean), if the proband was administered but could not complete an IQ test, indicated by the subject having a date for their IQ test but no IQ score, or if the subject had an HPO term or ICD code indicating intellectual disability or mental retardation. Age of walking unaided (in months) was taken from question 5A from the Autism Diagnostic Interview (ADI). We divided individuals into three possible categories for seizure status: yes, no, and unknown. A subject was put into the yes bin if he or she had a diagnosis of seizures or epilepsy, or a value of 2 on question 85 from the ADI (indicating a diagnosis of epilepsy). A subject was put into the no bin if no seizure/epilepsy diagnosis was indicated or if ADI question 85 had a value of 0. All remaining subjects were put into the unknown bin.

#### Burden of mutations in ASD as a function of IQ

*Burden of mutations over all genes*

We used full-scale IQ (FSIQ) to separate subjects into groups. Of the 5298 probands with any *de novo* mutation, 3010 have FSIQ information, 2055 (68.3%) with FSIQ > 70 and 1586 with FSIQ > 82. For a sample size N, the expected number of mutations within genes is computed as $E = 2N\phi$, where $\phi$ is the sum of the mutation rate, per variant type, over all relevant genes. (For example, to calculate $\phi$ for PTVs in genes with pLI > 0.995, we compute the sum of the PTV mutation rates for these genes.) We then compare $E$ to the observed count for this mutation class, O, and evaluate the distribution of O/E as a chi-square statistic with 1 degree of freedom.

*Burden of mutations over 102 TADA ASD genes*

This analysis addresses the question of whether the signal found in the 102 genes with q<0.10 could have arisen solely from low IQ subjects, such that any mutations found in higher IQ subjects occurred by chance. To answer this question, we must address the bias inherent in choosing 102 genes because they have q<0.10. To do so, we performed model-based simulations, similar to those used to evaluate the properties of the TADA model. We first select 874 genes with the smallest q-values from the real data and label them "signal genes". Let $M = 0.306N$ be the number of subjects with IQ < 70, who accumulate mutations at rates greater than chance. We generate mutations for the signal genes using the TADA model and Poisson rate ($2M\gamma\mu$), where $\mu$ is the gene-specific mutation rate and $\gamma$ is the increased rate of mutations due to this being a risk gene and the mutation of a particular type, and we generate additional mutations at a Poisson rate ($2[N - M]\mu$). We generate mutations in non-signal genes at a Poisson rate ($2N\mu$). We run TADA to get the new top 102 genes and the new signal genes, and we record counts occurring in new signal genes by chance (i.e., for individuals with high IQ). We perform the simulation 500 times to obtain the distribution of counts in signal genes for individuals with high IQ and compare this to the observed data. For all four informative mutation types, the expected counts were consistently lower than the observed count; only for missense mutations with MPC between 1 and 2 does the expected count, 13.54 (± 4.2) approach the observed value, 23 ($p=0.03$). For all other mutation types, the empirical p-value is far smaller, based on 500 simulations (MPC >2: 13.9 ± 3.9 versus 28; PTV for pLI >0.995: 8.3 ± 3.0 versus 48; and PTV for 0.5 < pLI < 0.995: 3.0 ± 1.9 versus 15). We also performed these simulations for a split on IQ at 82 and reached the same conclusion, that the mutations in the higher IQ ASD subjects accumulate at a rate far greater than chance.

#### Genes in recurrent genomic disorders

##### *Curation of reported GD loci*

We constructed a list of loci previously reported to be associated with ASD- or NDD-related phenotypes due to rare CNVs. We first collated coordinates of pathogenic genomic disorder (GD) regions as reported by nine previous studies (Sanders et al., 2015; Cooper et al., 2011; Wapner et al., 2012; Schaefer & Mendelsohn, 2013; Dittwald et al., 2013; Coe et al., 2014; Pinto et al., 2014; Wright et al., 2015; Rehm et al., 2015) and converted all coordinates to human reference genome build hg19 with UCSC liftOver tool as necessary. We next clustered the coordinates of all overlapping CNV regions using svtk bedcluster and a minimum 50% reciprocal overlap between segments, retaining the median clustered coordinates of all CNV regions appearing in at least two of the nine studies considered. After clustering, we excluded any CNV segments > 5Mb in size and all segments on sex chromosomes. Finally, we annotated each CNV segment passing all filters with all overlapping genes drawn from the list of autosomal genes considered during TADA analyses.

##### *Assessment of overlap between ASD-associated genes and GD loci*

We designed three permutation-based approaches to benchmark null expectations for the overlap of ASD-associated genes and GD loci. All approaches involved randomly drawing new sets of collinear genes for each GD locus from the list of all genes considered in TADA analyses, but differed in how these new genes were selected. These sampling approaches are summarized as follows:

1. Matched on number of genes: a new collinear list of genes was drawn for each GD locus, where the number of genes was matched to the number of genes in the original GD locus.
2. Matched on PTV mutation rates: a new collinear list of genes was drawn for each GD, where the number of genes was determined such that the sum of their estimated PTV mutation rates was at least as large as the sum of the estimated PTV mutation rates of the original list of genes in that GD locus.
3. Matched on brain expression, PTV mutation rates, and number of genes: prior to permutation, all genes were assigned a PTV mutation rate quintile and a brain expression quantile determined by the median brain expression value for that gene across all samples and all brain regions present in GTEx release v7 calculated after excluding genes with non-zero median brain expression. During permutation, a new collinear list of genes was drawn for each GD such that the number of genes matched the original GD locus, with the

additional requirements that the distribution of these genes across brain expression quintiles and PTV mutation rate quintiles were also preserved.

For each permutation, we performed one of the three above approaches for all 51 GD loci to obtain a new set of sampled genes, and we then counted the number of newly sampled genes that matched the TADA thresholds for ASD association in this study. We performed 1,000,000 permutations for each approach and computed p-values based on the fraction of all permutations where the number of GD loci with at least one randomly sampled ASD-associated gene matched or exceed the empirical observation in the original data. Fold-changes were determined as the observed number of GD loci with at least one ASD-associated gene divided by the mean number of GDs with at least one ASD-associated gene across all 1,000,000 permutations.

Finally, we titrated additional parameters to examine the variability of results from this permutation approach. For each of the three gene-sampling schemes above, we performed a separate 1,000,000 independent permutations for each combination of two additional factors, as follows:

1. ASD-associated gene list: we considered two different significance levels for ASD-associated genes, including (1) those determined as significant by the extended TADA model at FDR $\leq 0.1$ (n=102 genes) and (2) those reaching Bonferroni-corrected significance (n=26).
2. Chromosome sampling weights: for each GD locus in each permutation, an autosomal chromosome was selected based on one of two weighting schemes prior to randomly sampling a new set of collinear genes. These weights were either (1) determined by the fraction of all autosomal genes located on each chromosome, or (2) determined by the fraction of GD loci located on each chromosome.

All results were consistent across gene sampling strategies and the additional parameters had limited influence on our individual results or overall conclusions.

#### Enrichment of common variants in the detected genes

To investigate if the 102 ASD-associated genes were enriched for common variants associated with ASD and genetically correlated traits, we ran competitive gene-set enrichment analyses on the set of 102 genes using MAGMA (de Leeuw et al., 2015) and brain expressed genes from BrainSpan (see section below on developmental expression data) as background. We used summary statistics from the latest GWAS of ASD (Grove et al., 2017), ADHD (Demontis et al., 2017), major depression (Wray et al., 2018) (without the 23andMe contribution), schizophrenia (Ripke et al., 2014), and educational attainment (Lee et al. 2018), as well as a GWAS of height (Wood et al. 2014) as a control. In addition, to illustrate the effect of statistical power in the GWAS, we ran enrichment analyses also for historical GWAS for these phenotypes (Neale et al., 2010; Ripke et al., 2011; Rietveld et al., 2013; Ripke et al., 2013a; Ripke et al., 2013b; Okbay et al., 2016). From the summary statistics, gene-based p-values were estimated using test statistics defined as the sum of -log(SNP p-values) for SNPs located within the transcribed region plus a padding of 10kb flaking regions on either side. The gene-set enrichment was conducted by regressing gene-based Z-scores on a dichotomous gene-set indicator and covariates, which included gene size, gene density, sample size, the reciprocal of the minimal allele count, and the logarithm of these variables. We applied the default settings in MAGMA.

#### Expression analyses

*Tissue and CNS cell type-specific expression data*

Genotype-Tissue Expression (GTEx) RNA-seq data (https://www.gtexportal.org/home/) were summarized to GENCODE10 and gene-level reads per kilobase million mapped reads (RPKM) values were used across 53 tissue types, including 13 distinct brain regions. Samples with an RNA integrity number (RIN) $\leq 7$ were filtered and removed from subsequent analyses. Genes were defined as expressed if they were present at an RPKM of 0.5 in 80% of the samples from at least one tissue type, resulting in 27,546 transcripts. Finally, expression values were log-transformed ($\log_2$[RPKM+1]).

To determine tissue-specific gene expression signatures (*i.e.* genes which are significantly more expressed in a given tissue type compared to all other tissues), a linear regression model was applied for each gene for each tissue against all other tissues. Models were adjusted for age, RIN, gender, individual as a repeated measure, and surrogate variables to account for potential batch effects and other unwanted technical and biological variation. Significance values were adjusted for multiple testing using the Benjamini and Hochberg (BH) method to control the false discovery rate (FDR). After the BH correction, genes with Q-value < 0.05 and $\log_2$-fold change > 0.5 are defined as a genes exclusively expressed in a given tissue. These curated data formed the basis of our tissue-specific gene set enrichment analysis. All processed data are available upon request.

To test for over-representation of tissue and brain cell type-specific expression within a given gene set, a modified version of the GeneOverlap function in R was used so that all pairwise tests were corrected for multiple testing using the BH method. The Fisher's exact test function also provides an estimated odds ratio in comparison to a tissue- and genome-wide background set of 27,546 transcripts.

*Developmental expression data*

BrainSpan developmental RNA-seq data (www.brainspan.org) were summarized to GENCODE10 and gene-level RPKMs were used across 528 samples. Only the neocortical regions were used in our analysis—dorsolateral prefrontal cortex (DFC), ventrolateral prefrontal cortex (VFC), medial prefrontal cortex (MFC), orbitofrontal cortex (OFC), primary motor cortex (M1C), primary somatosensory cortex (S1C), primary association cortex (A1C), inferior parietal cortex (IPC), superior temporal cortex (STC), inferior temporal cortex (ITC), and primary visual cortex (V1C). Samples with RIN $\leq 7$ were filtered and removed from subsequent analyses. Genes were defined as expressed if they were present at an RPKM of 0.5 in 80% of the samples from at least one neocortical region at one major temporal epoch, resulting in 22,141 transcripts across 299 high-quality samples ranging from PCW 8 to 40 years of age (Figure S5). Finally, expression values were log-transformed ($\log_2$[RPKM+1]). All processed data are available upon request.

Linear regression was performed at each of 22,141 transcripts, modeling gene expression as a continuous dependent variable, as a function of a binary 'prenatal' stage variable. Similar to tissue-specific linear models (as above), each regression analysis included gender, individual as a repeated measure, ethnicity, and surrogate variables as adjustment variables. The regression model generated a 'prenatal effect' (*t*-statistic) of the $\log_2$-fold change of prenatal versus postnatal transcript abundance. A BH multiple test correction was used to control the FDR. Genes were defined as either prenatally or postnatally biased ($\log_2$ FC > 0.1 and q-value < 0.05) or unbiased in expression (q > 0.05). Under this paradigm, we measured the concordance of the prenatal effect across all 11 neocortical brain regions to ensure consistent gene-based effects and substantiate leveraging all 11 areas to represent one larger neocortical region (average $r = 0.956$). Subsequently, a total of five gene sets (ASD 102 genes, $ASD_P$ 53 genes, $ASD_{NDD}$ 49 genes, GER 58 genes, NC 24 genes) were evaluated by a Wilcoxon signed rank test to determine if the fetal effect distribution of the set differed significantly from the entire neocortical background, using the reduced statistic of one fetal effect per gene. The neocortical background was defined as genes which were simultaneously detected by WES in the current study as well as genes found to be expressed in the neocortex following quality control procedures (described above).

*Gene expression in cell types of the developing fetal neocortex*

The data selected from Nowakowski et al. (2017) consisted of 4,216 cells, which were divided into 25 cell types and expressed 58,865 identified transcripts. After matching and limiting transcripts to genes used for the TADA analysis, the expression data consisted of 17,116 genes, with the 102 TADA ASD genes included. Gene expression was first defined by developmental stage, in post-conception weeks (pcw). The 4,261 cells fell into 17 bins by developmental stage. For each developmental stage, a gene was expressed if at least one transcript mapped to this gene in 25% or more of cells for at least one pcw stage.

For cell type-specific analyses, six cell types were excluded because they could not be unambiguously associated with a cell type. The 19 remaining cell types contained 3,839 cells. Within each cell type cluster, a gene was considered expressed if one or more of its transcripts were detected in 25% or more cells, which resulted in 7,867 protein coding genes being expressed.

To determine enrichment for expression of 102 TADA ASD genes, the universe of genes expressed was determined by the cells in this experiment; e.g., for the cell type-specific enrichment, the universe was U=7,867 genes. The counts of the two-by-two table for enrichment of ASD genes, by cell type, were the number of TADA genes expressed in the cell type, X; Y, the number of TADA genes not expressed in the cell type; Z, the total number of genes expressed in the cell type minus X; and U – (X+Y+Z). Given this table, the enrichment odds ratio was calculated and significance judged by Fisher's exact test.

To evaluate how well cell types clustered, we first found 10% of genes showing the largest variation among cell types, as judged by analysis of variance. Performing hierarchical clustering using dissimilarity of expression for these genes revealed commonalities about the common types of cells (hclust package from R, centroid method).

To evaluate whether the enrichment odds ratio was a function of the number of genes expressed per cell type, we used linear regression. Because the odds ratio decreases as the number of genes increases, we next asked what might cause this phenomenon. It is reasonable to conjecture that the diversity of genes expressed (i.e., captured in our sequenced transcripts) would likely be a function of the evenness of expression of observed transcripts. In other words, if some transcripts were expressed at high levels, they will tend to lower the chance of capturing other genes in the sequence run for the cell, because the number of reads per cell was limited to fall between 1 and 2 million reads. This relative evenness should be captured by the mean expression over genes, and it is for these data: there is a very strong relationship between mean expression, taken over all expressed genes per cell type, and the number of genes expressed (R2 = 94.5%).

**Detecting Association With Networks (DAWN)**

The DAWN (Detecting Association With Networks) algorithm is a network-based gene discovery algorithm. The main assumption of the algorithm, related to this research, is that ASD-related genes are working as a functional group and should be co-expressed in a relevant spatio-temporal window during neurodevelopment. The algorithm predicts ASD-related genes using their interactions with other genes in the gene partial co-expression network, which is constructed using a Partial Neighborhood Selection method. A hidden Markov random field model assigns network-adjusted posterior risk scores to each gene (Liu et al., 2014; Liu et al., 2015). A gene is assigned a higher posterior risk score if the gene (1) has a high prior risk assigned using exome sequencing studies (i.e., TADA) and (2) is highly interacting with other risk genes.

To estimate the partial co-expression network, we used the BrainSpan microarray dataset (http://developinghumanbrain.org). We opted for this dataset over RNA-seq from an overlapping group of samples due to the larger number of samples and individuals in the microarray data. We extracted the subset of the data for a spatio-temporal window that was previously implicated for ASD risk: midfetal period and prefrontal cortex (PFC) (Willsey et al., 2013). Because the microarray probes for *CHD8*, one of the genes with the lowest p-values for ASD association (De Rubeis et al., 2014), do not perform well (Willsey et al., 2013; Cotney et al., 2015), we elected to impute the expression of *CHD8* in microarray data from the RNA-seq data. To do so, we picked the top genes that are highly correlated with *CHD8* in the RNA-seq data. Using the multivariate normal data imputation method, we imputed the corresponding measurements of *CHD8* in the microarray data for DAWN analysis. We input the TADA p-values as the prior risk scores. DAWN works with p-values less than, but not equal to, 1.0. Thus, we replaced the p-values of 7,994 genes (45.69%) with a p-value equal to 1.0 with 0.999. We fixed the top 10 ASD-associated genes with respect to TADA p-values as seed genes in the program. No additional covariates were supplied. The input hyperparameters for the DAWN algorithm were lambda = 0.24, p-value cutoff = 0.1, and partial correlation threshold = 0.7.

We used the DAWN results for a downstream network analysis of the interactions between synaptic genes and chromatin genes on a network of (i) ASD risk genes and (ii) additional genes chosen by DAWN as tightly interacting with ASD risk genes. The list of synaptic genes was obtained from Genes to Cognition (http://www.genes2cognition.org/db/GeneList). Specifically, we used lists L09-L16, which include the human orthologues of various synaptic complexes in mouse (1,123 genes). The list of chromatin modifiers was obtained from the Histome Database (Khare et al., 2012) and Huang et al. (2013) (173 genes total). We picked a threshold of FDR < 0.025, which yielded 100 genes in a DAWN ranking and evaluated interactions among them to form our subnetwork. Of these 100 genes, 40 genes have a TADA q-value < 0.1.

**Early developmental neocortical co-expression modules from tissue**

Weighted gene co-expression network analysis (WGCNA) was used to build a signed co-expression network from all early developmental samples passing our QC standards (as above). This resulted in 177 high-quality samples ranging from PCW 8 to 1 year of age that were used to build an early developmental network. The absolute values of the biweight midcorrelation coefficients were computed for all possible gene pairs and resulting values were transformed with an exponential weight (β) so that the final matrices followed an approximate scale-free topology (R2). We used a β threshold power of 21 so the subsequent network satisfied scale-free topology (R2>0.8), so the mean connectivity is high and the network contains enough information for module detection.

Module robustness was ensured by randomly sampling (2/3 of the total) from the initial set of samples 1000 times followed by consensus network analysis to define modules. The dynamic tree-cut algorithm was used to detect network modules with a minimum module size set to 200 and a cut tree height set to 0.9999. Singular value decomposition of each module's expression matrix was performed and the resulting module eigengene (ME), equivalent to the first principal component, was used to represent the overall expression profile for each module per sample. Pairs of modules were merged based on high correlation of ME values (>0.90). A total of 27 early developmental neocortical co-expression modules were identified. Each module was assessed for over-representation of five gene sets (ASD 102 genes, ASD enriched 53 genes, ID/DD/ASD 49 genes, GER 58 genes, NC 24 genes) using a one-sided Fisher's exact test and adjusting all pairwise tests for multiple testing using the BH method. Functional annotation of candidate modules was performed using ToppGene.

**Protein-protein connectivity among ASD candidate genes**

We used the InWeb_IM (Li et al., 2017) database of direct protein-protein interactions (PPI) to investigate whether the number of connections among $ASD_P$ and $ASD_{NDD}$ genes exceeds expectation, implicating significant functional relatedness. All candidate genes were brain-expressed as confirmed by the Allen Human Brain Atlas (http://portal.brain-map.org/) RNA-sequencing data, as this could serve a potential confounding factor for excessive connectivity compared to the general pool of genes present in the reference PPI network. We binned genes in each tested gene list into deciles of mean brain expression (in TPM) and used this distribution as a further reference for construction of random gene lists for significance analysis. To assess the significance of observed connectivity within each candidate gene list, we performed 1000 random draws of gene sets matching the candidate gene list in number of genes and distribution of expression levels. Empirical p-values were estimated as the proportion of random gene sets with greater or equal connectivity compared to the candidate gene list. Subsequently, to estimate the significance of connections with a known NDD gene list, we performed 1000 random draws of gene sets matching the candidate gene lists in number of genes and brain expression distribution. Further, the number of direct connections was estimated between a random gene set and NDD known genes. Empirical p-values were estimated as the proportion of random gene sets with a greater or equal number of connections to NDD known genes compared to the candidate gene list.

**Enrichment of transcription factor regulation network for the GER genes**

Starting with the 58 GER genes, we performed a systematic search for regulatory targets identified by protein-DNA interactions (e.g. ChIP-Seq) or protein-RNA interactions (e.g. iCLIP). First, we searched for target sets in available databases: ChEA (Lachmann et al., 2010) and ENCODE (ENCODE Project Consortium et al., 2007). The target lists were obtained from the Enrichr libraries (ChEA 2016 and ENCODE ChIP-seq 2015; Chen et al., 2013) and from a literature search for available ChIP or CLIP experiments, which are immunoprecipitation-based techniques to identify protein interactions with DNA and RNA, respectively. We found target data for 26 of 55 GER genes, with list of targets ranging from 5 to 8189 per gene. The data were generated in a wide range of tissues ranging from cell lines to liver to cortex and stemmed from 2 species (mouse and human). In total, we identified 21,514 gene targets from the 26 genes, which included 14,925 protein-coding genes from Ensembl. We constructed a network of 48,932 interactions with genes as the nodes and directed edges as transcription factor regulation relationships (Table S15).

To assess the significance of the connection between GER genes and a set of targets (i.e., NC genes), we generated random gene sets that matched genes with respect to brain expression, *de novo* PTV mutation rate, and pLI. For mutation rate, we used 5 bins such that each bin contained an equal number of genes (~3497). For pLI, we split genes into 3 groups: [0, 0.5), [0.5, 0.995) and [0.995, 1]. They contained 12152, 4506, and 1583 genes, respectively.

First, we checked whether GER genes are significantly connected to ASD risk genes. There are 409 connections between 58 GER genes and the 102 ASD risk genes (which contain the GER genes). Eight self-loops (a protein binding near to the gene that encodes it) were ignored. We generated 1,000 random gene sets of size 102. While the expected number of connections was 314, GER genes have 361 links to ASD risk genes ($p=0.006$). Then, we checked the significance of the connectivity within GER genes (only GER-GER connections). There are 229 connections within GER genes. Repeating the same analysis, we found the expected number of connections to be 175, demonstrating significant connectivity ($p < 0.001$). On the other hand, the connectivity between GER genes and NC genes was weaker—there are 132 connections, whereas the expected was 140 ($p=0.72$).

As a further control, we checked whether GER genes were significantly connected to congenital heart disease risk genes, which is known to have an overlapping genetic component with ASD (Jin et al., 2017). The list of 253 congenital heart disease risk genes was obtained from Jin et al. (2017). Of the 253 genes, 19 were excluded because they are either not autosomal protein-coding genes and/or lack known mutations rates, as used in our analysis. Specifically, 11 are on chromosome X (*HCCS, NSDHL, RBM10, PQBP1, ZIC3, GPC3, BCOR, FLNA, OFD1, COX7B, MID1*); 6 have no data in ExAC (*DNAH11, C1ORF127, IRX5, GATA6, FOXC2, FOXC1*); and 2 are noncoding: (*RNU4ATAC* and *RPS17*). There are 685 links between GER genes and 234 congenital heart disease genes, and this number has a p-value of 0.007 (expected 622; genes were matched with respect to pLI and mutation rate but not brain expression). Thus, despite a strong regulation relationship existing within GER genes and even between GER genes and congenital heart disease genes, this is missing between GER genes and NC genes, suggesting that these functional circuitries act independently rather than as a coherent unit, which is also seen in our DAWN analysis.

**Enrichment of CHD8 targets for the GER genes**

To assess whether ASD risk genes relate to known genome-wide CHD8 binding sites, we tested our previously defined GER and NC gene sets for enrichment with human brain-specific sequences from two independent ChIP sequencing (ChIP-seq) studies covering: 1) 3,281 CHD8-binding sites in the human mid-fetal brain at 16-19 pcw (Cotney et al., 2015); and 2) 6,860 CHD8-binding sites in human neural progenitor cells (NPCs), which reflects the intersection of signal-enriched regions detected by all three CHD8 antibodies used in the study (Sugathan et al., 2014). In order to assess overlap with these binding sites, genomic coordinates were defined as the start and end positions for each GER and NC gene (analogous to gene length). A permutation-based approach with 1,000 random permutations was used to determine statistical significance of the overlap between genomic coordinates for GER and NC genes with CHD8-binding sites using the R package regioneR (Gel et al., 2015).

#### DATA AND SOFTWARE AVAILABILITY

All data generated as part of the ASC is transferred to dbGaP with Study Accession: phs000298.v4.p3. TADA has been previously described (He et al., 2013) and enhancements to TADA are described in detail in the main text and the STAR Methods. DAWN has also been described (Liu et al., 2014).

**KEY RESOURCES TABLE**

| REAGENT or RESOURCE | SOURCE | IDENTIFIER |
|---|---|---|
| **Deposited Data** | | |
| ASC-generated WES sequencing data | This paper | dbGaP Study Accession: phs000298.v4.p3 |
| Human reference genome NCBI build 37, GRCh37 | Genome Reference Consortium | http://www.ncbi.nlm.nih.gov/projects/genome/assembly/grc/human/ |
| Exome aggregation consortium (ExAC) | Let et al., 2016 | http://exac.broadinstitute.org/ |
| Genome aggregation database (gnomAD) | Karczewski et al., 2019 | https://gnomad.broadinstitute.org/ |
| Deciphering Developmental Disorders (DDD) | DDD, 2017 | https://www.ddduk.org/ |
| Genotype-Tissue Expression (GTEx) resource | GTEx Consortium, 2017 | https://gtexportal.org/home/ |
| BrainSpan | Li et al., 2018 | http://www.brainspan.org/ |
| Single-cell RNA-seq data from developing cortex | Nowakowski et al., 2017 | NA |
| InWeb_IM (protein-protein interaction data) | Li et al., 2017 | http://www.lagelab.org/resources/ |
| **Software and Algorithms** | | |
| Genome Analysis Toolkit (GATK) | Van der Auwera et al., 2013 | https://software.broadinstitute.org/gatk/ |
| Hail | https://hail.is/ | https://github.com/hail-is/hail/ |
| Variant Effect Predictor (VEP) | McLaren et al., 2016 | http://grch37.ensembl.org/Homo_sapiens/Tools/VEP |
| TADA | He et al., 2013 | http://www.compgen.pitt.edu/TADA/TADA_guide.html |
| Gene Ontology (via Panther) | Mi et al., 2019 | http://www.pantherdb.org/ |
